# Enhancer-associated H3K4 methylation safeguards in vitro germline competence

**DOI:** 10.1101/2020.07.07.192427

**Authors:** Tore Bleckwehl, Giuliano Crispatzu, Kaitlin Schaaf, Patricia Respuela, Michaela Bartusel, Laura Benson, Stephen J. Clark, Kristel M. Dorighi, Antonio Barral, Magdalena Laugsch, Wilfred F. J. van IJcken, Miguel Manzanares, Joanna Wysocka, Wolf Reik, Álvaro Rada-Iglesias

## Abstract

Germline specification in mammals occurs through an inductive process whereby competent cells in the post-implantation epiblast differentiate into primordial germ cells (PGC). The intrinsic factors that endow epiblast cells with the competence to respond to germline inductive signals remain unknown. Single-cell RNA sequencing across multiple stages of an *in vitro* PGC-like cells (PGCLC) differentiation system shows that PGCLC genes initially expressed in the naïve pluripotent stage become homogeneously dismantled in germline competent epiblast like-cells (EpiLC). In contrast, the decommissioning of enhancers associated with these germline genes is incomplete. Namely, a subset of these enhancers partly retain H3K4me1, accumulate less heterochromatic marks and remain accessible and responsive to transcriptional activators. Subsequently, as *in vitro* germline competence is lost, these enhancers get further decommissioned and lose their responsiveness to transcriptional activators. Importantly, using H3K4me1 deficient cells, we show that the loss of this histone modification reduces the germline competence of EpiLC and decreases PGCLC differentiation efficiency. Our work suggests that, although H3K4me1 might not be essential for enhancer function, it can facilitate the (re)activation of enhancers and the establishment of gene expression programs during specific developmental transitions.

## INTRODUCTION

Competence can be defined as the ability of a cell to differentiate towards a specific cell fate in response to intrinsic and extrinsic signals^1^. While the extracellular signals involved in the induction of multiple cell fates have been described^2^, the intrinsic factors that determine the cellular competence to respond to those signals remain elusive. One major example illustrating the dynamic and transient nature of competence occurs early during mammalian embryogenesis, as the primordial germ cells (PGC), the precursors of the gametes, become specified. In mice, following implantation and the exit of naïve pluripotency (E4.5 - E5.5), PGC are induced from the epiblast around E6.0 ∼ E6.25 at the proximo-posterior end of the mouse embryo^3^. The induction of PGC occurs in response to signals emanating from the extraembryonic tissues surrounding the epiblast: BMP4 from the extraembryonic ectoderm and WNT3 from the visceral endoderm^4^. Furthermore, regardless of their position within the embryo, formative epiblast cells (∼E5.5-6.25)^5–7^ are germline competent when exposed to appropriate signals, but this ability gets lost as the epiblast progresses towards a primed pluripotency stage (>E6.5)^4^. However, the intrinsic factors that confer germline competence on the formative but not the primed epiblast cells remain obscure^8, 9^, partly due the limited cell numbers that can be obtained *in vivo* from mouse peri-implantation embryos (E4.5-E6.5). These limitations were mitigated by a robust *in vitro* differentiation system whereby mouse embryonic stem cells (ESC) grown under 2i conditions (naïve pluripotency) can be sequentially differentiated into EpiLC and PGCLC that resemble the formative epiblast and E9.5 PGC, respectively^10^. This system facilitated the mechanistic and genomic characterization of transcription factors (TFs)^11–13^ and epigenetic reprogramming events^14–16^ previously shown to be involved in PGC specification *in vivo*^17, 18^.

Using the PGCLC *in vitro* differentiation system, two transcription factors, FOXD3 and OTX2, were found to promote the transition from naïve to formative pluripotency by coordinating the silencing of naïve genes and the activation of early post-implantation epiblast markers^9, 19, 20^. Subsequently, FOXD3 and OTX2 restrict the differentiation of EpiLC into PGCLC and, thus, the silencing of these TFs is required for *in vitro* germline specification^9, 20^. The regulatory function of FOXD3 during these developmental transitions involves binding to and silencing of enhancers shared between naïve pluripotent cells and PGCLC^9^. Interestingly, during the transition from 2i ESC to EpiLC, FOXD3-bound enhancers lose TF and co-activator binding as well as H3K27ac but partly retain H3K4me1. This suggests that these enhancers do not become fully decommissioned, but transiently display a chromatin state similar, but not identical, to that of primed enhancers^21, 22^. Enhancer priming typically involves binding of pioneer TFs and pre-marking by H3K4me1 that can precede and facilitate subsequent enhancer activation (*i.e.* marking by H3K27ac, recruitment of RNA Pol II, production of eRNAs)^23–26^. Interestingly, in differentiated macrophages, enhancers activated upon stimulation rapidly lose H3K27ac and TF binding, while retaining H3K4me1 for considerably longer. It was proposed that H3K4me1 persistence could facilitate a faster and stronger enhancer induction upon restimulation^27^. It is currently unknown whether, during development, H3K4me1 persistence once enhancers become decommissioned can similarly facilitate their eventual re-activation^21^.

H3K4me1 is catalyzed by the SET domains of the histone methyltransferases MLL3 (KMT2C) and MLL4 (KMT2D), which are part of the COMPASS Complex^28, 29^. The knockout (KO) of *Mll3/4* or *Mll4* alone impairs enhancer activation and results in differentiation defects in various lineages^30–37^. However, the importance of MLL3/4 for enhancer function might be independent of H3K4me1 deposition, since *Mll3/4* catalytic mutant ESC (*Mll3/4* dCD) in which H3K4me1 was globally lost, only displayed minor defects in enhancer activation (*i.e.* H3K27ac, RNA Pol II and eRNA levels) in comparison to *Mll3/4* KO ESC^38^. Similarly, work in *Drosophila melanogaster* showed that, while the KO of *Trr*, the homolog of *Mll3/4* in flies, was embryonic lethal, an amino acid substitution in the SET domain of *Trr* that globally reduced H3K4me1 did not impair development or viability^39^. On the other hand, subsequent work with *Mll3/4* dCD ESC showed that the recruitment of chromatin remodelers^40^ and the establishment of long-range chromatin interactions^32^ required H3K4me1. Hence, the functional relevance of H3K4me1 for enhancer function is still under debate^41^.

Here we performed an extensive transcriptional and epigenetic characterization of the main stages of PGCLC differentiation to gain insights into the molecular basis of germline competence. Comparisons between germline competent EpiLC and non-competent EpiSC revealed that a notable fraction of PGCLC enhancers, which tend to be already active in the naïve stage (*i.e.* 2i ESC), partly retained H3K4me1 and remained accessible and responsive to TFs in EpiLC in comparison to EpiSC. Most importantly, the persistence of H3K4me1 within PGCLC enhancers seems to contribute to *in vitro* germline competence, as in the absence of this histone mark, PGCLC differentiation efficiency is reduced.

## RESULTS

### Characterization of the PGCLC *in vitro* differentiation system by single-cell RNA-seq

To overcome the scarcity and transient nature of PGCs *in vivo,* we used the *in vitro* PGCLC differentiation system^10^. Thereby mouse ESC are differentiated into EpiLC from which PGCLC can be obtained within heterogeneous embryoid bodies (EB). In contrast, Epiblast stem cells (EpiSC), resembling the post-implantation gastrulating epiblast, cannot be efficiently differentiated into PGCLC and, thus, display limited germline competence (Fig. 1a)^7, 10^. To better characterize this *in vitro* system, we performed single-cell RNA sequencing (scRNA-seq) across multiple stages of PGCLC differentiation (the scRNA-seq data can be easily explored with the *cloupe* file available through GEO: GSE155088). t-SNE analysis of the resulting single cell transcriptomes (Supplementary Data 1) showed that cells tend to cluster within their corresponding differentiation stage (Fig. 1b). However, Day 2 and Day 4 EB showed cellular heterogeneity and formed distinct subclusters (Fig. 1b-c). One of these subclusters was identified as PGCLC based on the high expression of major PGC markers (e.g. *Prdm14*, *Prdm1, Tfap2c, Dppa3*), while the additional subpopulations within d4 EB were annotated based on the expression of cell identity markers identified by single-cell transcriptional profiling of E8.25 mouse embryos^42^ (Fig. 1c, Supplementary Fig. 1a-c). Remarkably, these subclusters were similar to the extraembryonic tissues (*i.e.* extraembryonic ectoderm, extraembryonic mesoderm and endothelium) that surround PGCs in the proximo-posterior end of the mouse embryo following germline specification *in vivo*. Furthermore, differential expression analysis between the PGCLC cluster and the remaining cells of the d2 and d4 EBs (see Methods) led to the identification of a set of 389 PGCLC genes (Supplementary Data 2), which included the PGC markers mentioned above as well as major naïve pluripotency regulators (e.g. *Nanog*, *Esrrb*) (Fig. 1d-e). In agreement with previous reports^10^, we found that several PGCLC genes were highly expressed in ESC, progressively silenced in EpiLC and finally reactivated in PGCLC (Fig. 1d-e, Supplementary Fig. 1d). Lastly, we defined gene sets specific for the three investigated *in vitro* pluripotent cell types (i.e. ESC, d2 EpiLC and EpiSC) by performing differential gene expression analysis between each cell type and the other two (Supplementary Data 2).

**Fig. 1:**
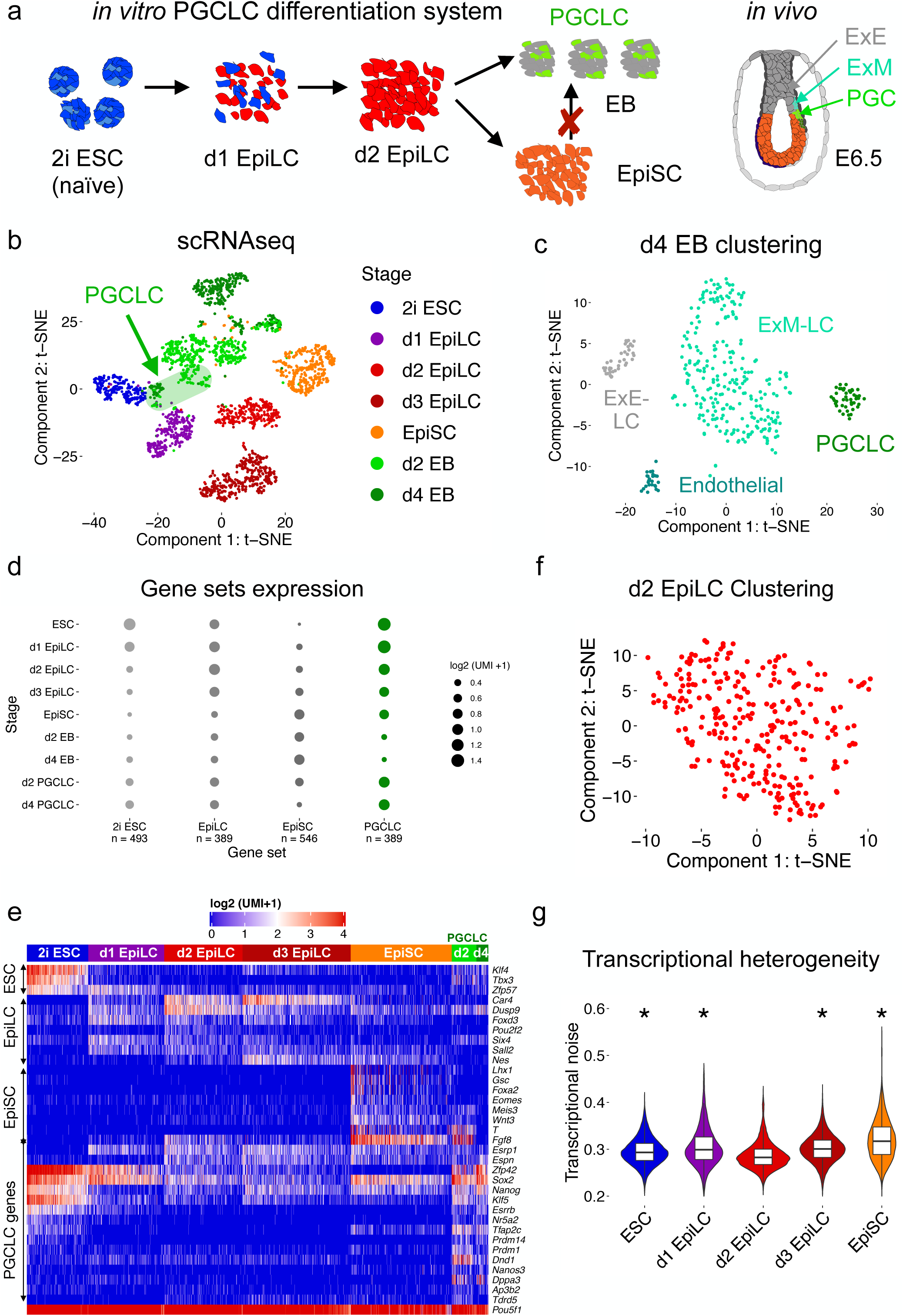
Single-cell RNA-sequencing (scRNA-seq) profiling of the *in vitro* PGCLC differentiation system. a.) Schematic representation of the *in vitro* PGCLC differentiation system. Embryonic stem cells (ESC) are differentiated into epiblast-like cells (EpiLC). Day 2 (d2) EpiLC are germline competent and can be differentiated further into embryoid bodies (EB) containing primordial germ cell-like cells (PGCLC). In contrast, further progress of EpiLC into epiblast stem cells (EpiSC) restricts germline competence. The diagram to the far right illustrates the *in vivo* location of PGCs at the proximal-posterior end of a mouse embryo. b.) t-SNE plot of the scRNA-seq data (n = 2,782 cells) generated across the main stages of the *in vitro* PGCLC differentiation protocol. The PGCLC identified within the d2 and d4 EB are shadowed in green. c.) t-SNE plot based on the scRNA-seq data generated in d4 EB (n = 368 cells). K-means clustering of d4 EB identified four clusters resembling the transcriptomes of PGCs and their main surrounding tissues *in vivo*: Extraembryonic Ectoderm (ExE), Extraembryonic Mesoderm (ExM) and Endothelial cells. d.) Gene expression summary for the different gene sets defined using the single-cell RNA-seq data. The ESC, EpiLC and EpiSC gene sets were determined by considering the genes differentially expressed in each stage compared to the remaining two (i.e. ESC vs. d2 EpiLC and EpiSC). The PGCLC gene set was determined by considering the differentially expressed genes in the d2 and d4 PGCLC cluster compared to the remaining cells of the d2 and d4 EBs. The number of genes per set are indicated below and the size of a dot represents the average expression of all genes within a gene set for the indicated cellular stages. e.) Heatmap showing the expression of selected genes from the gene sets shown to the left (i.e. ESC, EpiLC, EpiSC and PGCLC gene sets) within individual cells belonging to the indicated cellular stages (colored above the heatmap). The expression values are displayed as UMI (unique molecular identifier) counts. *Pou5f1*, which is not assigned to any specific gene set as it is similarly expressed in all cellular stages, is shown to illustrate the quality of the single-cell RNA-seq data. f.) t-SNE plot for the d2 EpiLC scRNA-seq data alone (n = 289 cells), without any obvious clusters. g.) Violin plots showing transcriptional noise, defined as cell-to-cell transcript variation for the 500 most variable genes, for ESC, EpiLC and EpiSC. Lower transcriptional noise indicates higher transcriptional similarity between the cells belonging to a particular stage. All cellular stages shown were compared to d2 EpiLC using Wilcoxon tests (*: p-value < 2.2 10^−16^).

### Germline competent EpiLC are transcriptionally homogeneous

Previous work indicates that the acquisition of germline competence in day 2 (d2) EpiLC entails the complete dismantling of the naïve gene expression program^5, 6^, which is then partly re-activated during PGCLC induction. In agreement with this, all d2 EpiLC clustered together and neither a distinct subpopulation indicative of a retained naïve pluripotency expression program nor signs of precocious germline induction could be identified (Fig. 1b,f; Supplementary Fig. 1e). Congruently, the cell-to-cell variability in gene expression levels, defined as transcriptional noise, was significantly lower in d2 EpiLC than in the preceding or subsequent cellular stages (Fig. 1g). This is in agreement with the transcriptional homogeneity of the E5.5 epiblast *in vivo*^43^ (Supplementary Fig. 1f), which bears the lowest transcriptional noise during mouse peri-implantation development^44^. Lastly, we analyzed bulk RNA-seq data generated in *Otx2^−/−^*^20^ and *Prdm14^−/−^*^14^ EpiLC, which despite displaying increased and decreased germline competence, respectively, showed normal expression of PGCLC genes (Supplementary Fig. 1g). Therefore, in agreement with previous work^6^, our scRNA-seq analysis indicates that *in vitro* germline competence entails a transcriptionally homogeneous stage in which the gene expression program shared between naïve pluripotency and PGCLC is silenced.

### Identification of PGCLC active enhancers

Many PGCLC genes, especially those active in ESC, are silenced in EpiLC and EpiSC (Fig. 1d,e, Supplementary Fig. 1d), yet only EpiLC display high germline competence. Taking previous observations into account^9, 45, 46^, we hypothesized that enhancers involved in the induction of PGCLC genes might display epigenetic differences between EpiLC and EpiSC that could explain their distinct germline competence. To test this hypothesis, we first identified distal H3K27ac peaks in d6-sorted PGCLC using publically available data^15^. In agreement with our previous observations^9^, a large fraction of the d6 PGCLC H3K27ac peaks were initially active in ESC, lost H3K27ac in EpiLC and became progressively reactivated in d2 and d6 PGCLC (Supplementary Fig. 2a). Since most of the d6 PGCLC H3K27ac peaks were also active in ESC, we then used Capture Hi-C data generated in ESC^47^ to systematically associate these distal peaks to their putative target genes. Finally, we defined PGCLC enhancers as those distal d6 PGCLC H3K27ac peaks that could be physically linked to our PGCLC gene set (Supplementary Data 2). This resulted in 415 PGCLC enhancers linked to 216 of the 389 PGCLC genes (Fig. 2a). Furthermore, to compare epigenetic changes between different enhancer groups, EpiLC and EpiSC enhancers were defined using similar criteria (Methods; Supplementary Data 3).

**Fig. 2:**
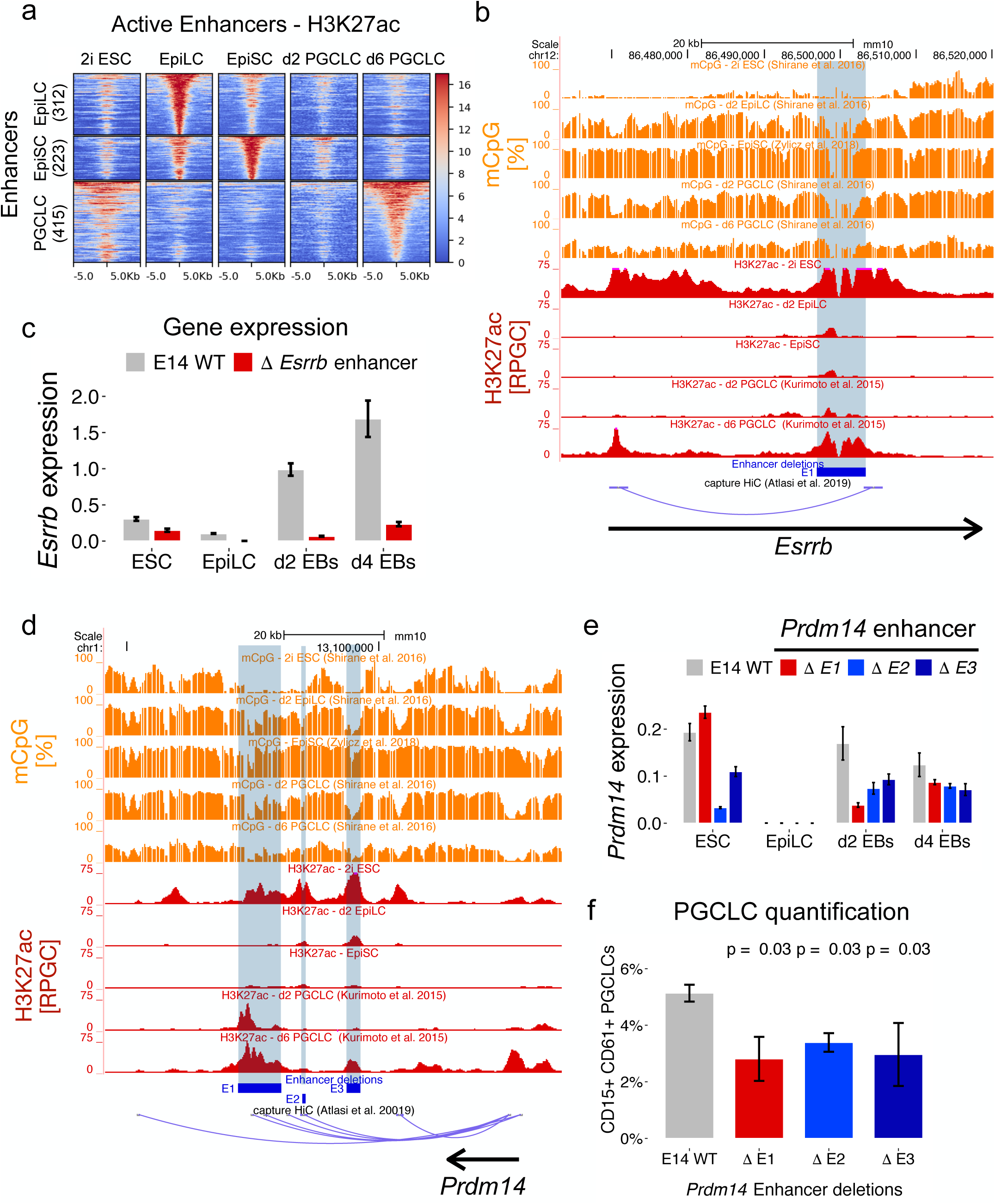
Identification and functional assessment of PGCLC enhancers. a.) Heatmaps showing the H3K27ac dynamics for EpiLC, EpiSC and PGCLC enhancers during PGCLC differentiation. For EpiLC: 146 out of 389, EpiSC: 164 out of 546 and PGCLC: 216 out of 389 genes from the gene sets defined in Fig. 1 were linked to their enhancers (see *Methods* for the criteria used to define the different enhancer sets). Enhancers within each set were ordered according to the H3K27ac levels in the corresponding cell type. b.) Genome-browser view of the H3K27ac and CpG methylation dynamics during PGCLC differentiation for a PGCLC enhancer, highlighted in blue, found within the *Esrrb* locus. The enhancer is physically linked to the *Esrrb* transcription start site (TSS) in ESC according to capture Hi-C data from Atlasi *et al.* 2019. The blue rectangle denotes the enhancer deletion generated in mESC using CRISPR/Cas9 technology. c.) *Esrrb* expression levels were measured by RT-qPCR in d4 EB differentiated from WT ESC and the two different ESC clonal lines with the *Esrrb* enhancer deletion shown in (b). The expression values were normalized to two housekeeping genes (*Eef1a1* and *Hprt*). Error bars represent standard deviations from 6 measurements (two clonal lines x three technical replicates). d.) Genome-browser of the H3K27ac and CpG methylation dynamics during PGCLC differentiation for three PGCLC enhancers (E1-E3) found within the *Prdm14* locus. The blue rectangles denote the enhancer deletions generated in mESC using CRISPR/Cas9 technology. The E1-E2 enhancers are physically linked to the *Prdm14* TSS in ESC according to capture Hi-C data from Atlasi *et al.* 2019. e.) *Prdm14* expression levels were measured by RT-qPCR in d4 EB differentiated from WT ESC and ESC with the indicated *Prdm14* enhancer deletions (two clonal lines for each deletion). The expression values were normalized to two housekeeping genes (*Eef1a1* and *Hprt*). Error bars represent standard deviations from 6 measurements (two clonal lines × three technical replicates). f.) WT ESC and ESC lines with the indicated *Prdm14* enhancer deletions were differentiated into PGCLC. PGCLC were measured as CD15^+^CD61^+^ cells within d4 EB. Each PGCLC quantification was performed in biological duplicates and two different clonal lines were used for each enhancer deletion (n=2×2). The percentages of PGCLC obtained when differentiating the ESC with the enhancer deletions were compared to those obtained with WT ESC using Wilcoxon tests.

To validate our PGCLC enhancer calling strategy, we first selected three representative enhancers linked to *Esrrb*, *Klf5* and *Lrrc31/Lrrc34*, respectively (Fig. 2b, Supplementary Fig. 2b). These three enhancers were initially active in ESC, got silenced in EpiLC and finally became re-activated in PGCLC. Each enhancer was individually deleted in ESC using CRISPR/Cas9 technology (Supplementary Fig. 2c). The deletion of the enhancers associated with *Lrrc31/Lrrc34* and *Klf5* significantly reduced the expression of the corresponding target genes in ESC and d4 EB (Supplementary Fig. 2d-e). The *Esrrb* enhancer deletion had a moderate effect in ESC and severely diminished *Esrrb* expression in d4 EB (Fig. 2c). Since some PGCLC genes are associated with multiple and potentially redundant regulatory elements (*i.e.* 415 enhancers linked to 216 PGCLC genes; 1.9 enhancers/gene), we individually deleted three different enhancers (*i.e.* E1, E2 and E3) (Fig. 2d; Supplementary Fig. 2c) previously described as components of a *Prdm14* super-enhancer^48^. In agreement with previous work, the E2 deletion strongly reduced *Prdm14* expression in ESC, while the deletion of E3 or E1 had considerably smaller effects (Fig. 2e)^48^. Upon PGCLC differentiation, the regulatory importance of each enhancer changed and E1 clearly contributed to *Prdm14* expression, especially during early PGCLC induction (Fig. 2e). These results suggest that, rather than being components of an ESC super-enhancer, the E1-E3 elements differentially contribute to *Prdm14* expression in either ESC (*i.e.* E2) or PGCLC (*i.e.* E1). Furthermore, in agreement with the role of *Prdm14* as a PGC master regulator^49^, we found that the individual E1-E3 deletions significantly impaired PGCLC differentiation (Fig. 2f, Supplementary Fig. 2f). Altogether, the previous deletions support the relevance of the identified PGCLC enhancers and suggest that some of them (e.g. *Esrrb* and *Prdm14* E1 enhancers) are particularly relevant during PGCLC induction, while others might be important in both ESC and PGCLC.

### Partial decommissioning of PGCLC enhancers in EpiLC

To explore whether PGCLC enhancers display any chromatin differences between EpiLC and EpiSC, we generated an extensive set of ChIP-seq and ATAC-seq data sets in ESC, EpiLC and EpiSC (Fig. 3a; Supplementary Fig. 3a). Overall, the most pronounced epigenetic changes within PGCLC enhancers were observed for H3K4me1, which was higher in EpiLC, and H3K9me3 and mCpG, which were increased in EpiSC (Fig. 3a-b). These differences were not observed for other enhancer groups (e.g. EpiSC enhancers), suggesting that they were not due to technical reasons or global epigenomic differences between EpiLC and EpiSC (Fig. 3a). Moreover, when analyzing the transcription start sites (TSS) of the PGCLC genes we found that, although H3K4me1 was higher in EpiLC than in EpiSC, its overall levels were low compared to PGCLC enhancers (Fig. 3a). Similarly, constitutive heterochromatin marks (e.g. H3K9me3, mCpG) around TSS increased in EpiSC, but their levels were lower than within PGCLC enhancers. Other chromatin features typically found at promoter regions (e.g. H3K4me2/3, high chromatin accessibility) were similar around PGCLC TSS in EpiLC and EpiSC (Fig. 3a). Therefore, subsequent analyses were focused on PGCLC enhancers rather than promoters.

**Fig. 3:**
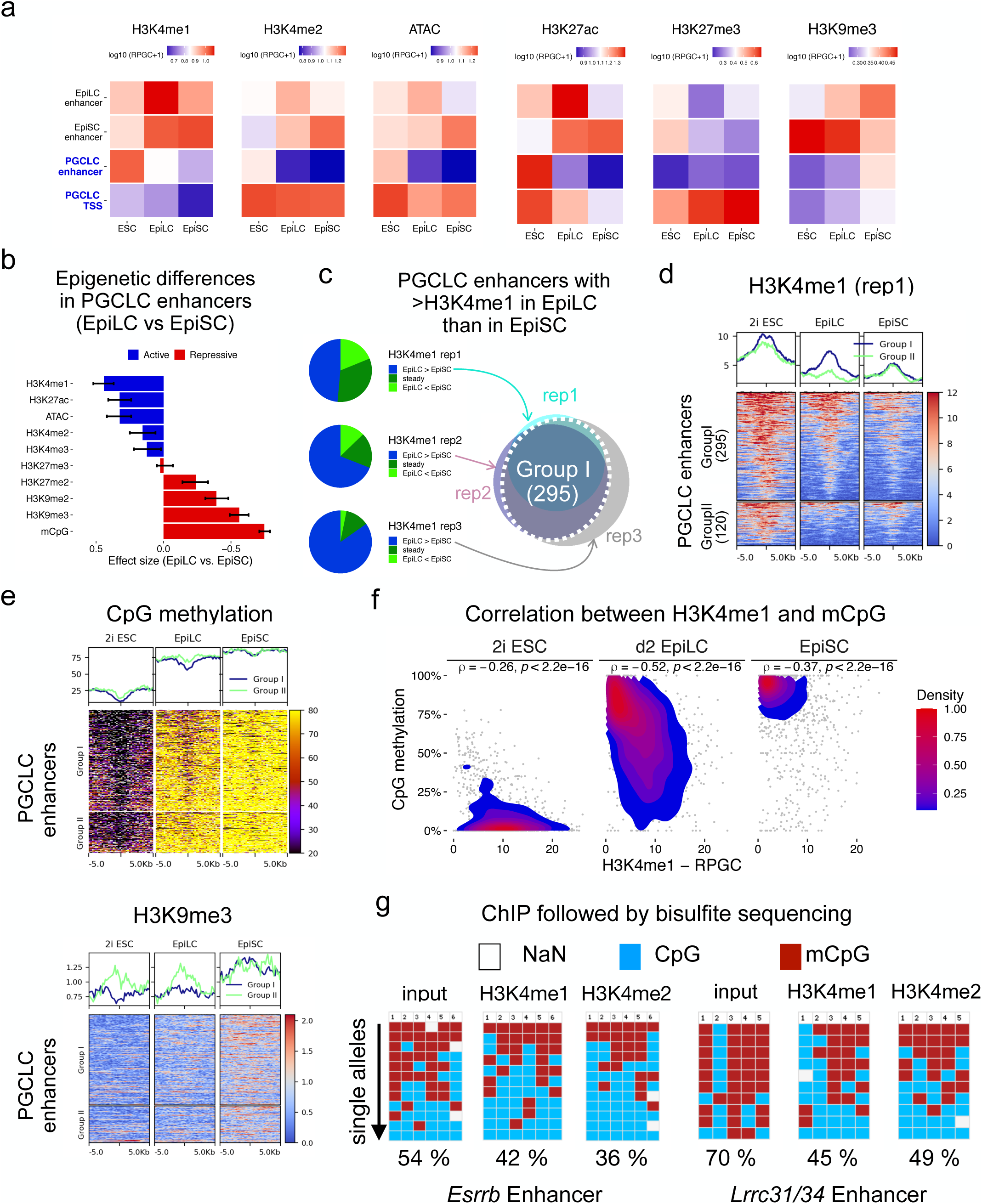
Evaluation of the epigenetic features of PGCLC enhancers in ESC, EpiLC and EpiSC. a.) Summary plot showing the average levels of the different epigenetic marks measured in 2i ESC, d2 EpiLC and EpiSC for the EpiLC, EpiSC and PGCLC enhancers as well as the TSS of the PGCLC genes. Quantifications were performed by measuring the average signals of each epigenetic mark within −/+ 1kb of the enhancers orTSS. RPGC: Reads per genomic content. b.) The magnitude of the differences between EpiLC and EpiSC when comparing the levels of the different epigenetic marks within PGCLC enhancers was determined as the effect sizes from paired Wilcoxon tests. The error bars represent the confidence intervals. c.) Three H3K4me1 ChIP-seq replicates were generated in d2 EpiLC and EpiSC, respectively. For each replicate, the average H3K4me1 signal within −/+ 1kb of each PGCLC enhancer was calculated and compared between EpiLC and EpiSC. Those PGCLC enhancers showing an EpiLC/EpiSC ratio higher than 1.2-fold were assigned to the ‘EpiLC > EpiSC’ category for each replicate. PGCLC enhancers assigned to the ‘EpiLC > EpiSC’ category in at least two replicates were defined as *Group I* (Venn diagram). All other enhancers, including those from the ‘steady’ (EpilC/EpiSC H3K4me1 ratio: 0.8>FC>1.2) and ‘EpiLC < EpiSC’ (EpiLC/ EpiSC H3K4me1 ratio: FC<0.8) categories, were assigned to *Group II*. d.) Average profile (top) and heatmap (bottom) plots showing H3K4me1 levels for the Group I and Group II PGCLC enhancers in 2i ESC, d2 EpiLC and EpiSC. The H3K4me1 signals correspond to one of the ChIP-seq replicates from the E14Tg2 ESC cell line. e.) Average profile (top) and heatmap (bottom) plots showing H3K9me3 levels and the percentage of CpG methylation for the Group I and Group II PGCLC enhancers in 2i ESC, d2 EpiLC and EpiSC. f.) Correlation plots between mCpG methylation and H3K4me1 at Group I PGCLC enhancers in 2i ESC, d2 EpiLC and EpiSC. For each enhancer the average percentage of CpG methylation or level of H3K4me1 was calculated for two 500 bp bins up- and downstream of the enhancer. All four bins of one enhancer are plotted and the spearman correlation was determined. Shown are the measurements of the first H3K4me1 ChIP-seq replicate. g.) The CpG methylation status of the PGCLC enhancers associated with *Esrrb* and *Lrrc31/34* was investigated by bisulfite sequencing using as templates ChIP input DNA, H3K4me1 ChIP DNA and H3K4me2 ChIP DNA generated in d2 EpiLC. The columns of the plots correspond to individual CpG dinucleotides located within each enhancer. Unmethylated CpGs are shown in blue, methylated CpGs in red and CpGs that were not sequenced are shown in gray. At least 10 alleles were analyzed for each template DNA (rows). In (e) and (f) the CpG methylation data was obtained from Habibi *et al.* 2013 and Zylicz *et al.* 2015.

To identify PGCLC enhancers showing consistent differences between EpiLC and EpiSC, we focused on H3K4me1, which is widely considered as an enhancer mark^50^. We generated two additional H3K4me1 ChIP-seq replicates in EpiLC and EpiSC (including data from a different ESC strain *i.e.* R1) and determined that 71% of the PGCLC enhancers showed higher H3K4me1 levels in EpiLC than in EpiSC in at least two replicates (71% Group I; 29% Group II) (Fig. 3c-d; Supplementary Fig. 3b-c). The Group I PGCLC enhancers also retained more H3K27ac in EpiLC, while the differences in H3K4me2 and chromatin accessibility, as measured by ATAC-seq, were rather moderate (Supplementary Fig. 3d). In contrast, the Group I enhancers displayed lower mCpG and H3K9me3 levels in EpiLC than in EpiSC, while these differences were considerably less obvious for the Group II enhancers (Fig. 3d-e, Supplementary Fig. 3e). Furthermore, H3K4me1 and mCpG were inversely correlated across Group I PGCLC enhancers, particularly in EpiLC (Fig. 3f). This negative correlation was also investigated by performing H3K4me1 ChIP in EpiLC followed by bisulfite sequencing of two representative Group I PGCLC enhancers (*i.e. Esrrb* and *Lrrc31/Lrrc34* enhancers) (Fig. 3g). In comparison to the input genomic DNA, the H3K4me1-enriched DNA showed lower mCpG levels at the two analyzed enhancers (Fig. 3g), further supporting the antagonism of CpG methylation and H3K4 methylation^51, 52^.

In summary, a subset of the PGCLC enhancers show partial retention of H3K4me1 and lower levels of heterochromatic marks (i.e. mCpG, H3K9me3) in EpiLC compared to EpiSC (Fig. 3d-e). These enhancers tend to be initially active in ESC but lose H3K27ac and TF/coactivator binding (indirectly measured by ATAC-seq) already in EpiLC (Supplementary Fig. 3d). Therefore, our data suggest that a significant fraction of PGCLC enhancers are not fully decommissioned in EpiLC, which we hypothesize could facilitate their re-activation in PGCLC and, thus, contribute to germline competence.

### Impaired decommissioning of PGCLC enhancers in *Otx2* deficient cells

The deletion of *Otx2* increases and prolongs germline competence in EpiLC^20^. Hence, to further explore the relationship between the chromatin features of PGCLC enhancers and germline competence, we analyzed *Otx2^−/−^* cells ^53^. Firstly, we confirmed the extended germline competence of *Otx2^−/−^* cells, which can be robustly differentiated into PGCLC after keeping them for up to seven days in EpiSC culture conditions (Fig. 4a, Supplementary Fig. 4a). ChIP-seq experiments revealed that the increased germline competence of *Otx2^−/−^* cells was correlated with the retention of H3K4me1, H3K4me2 and H3K27ac in PGCLC enhancers, particularly within those displaying partial decommissioning in WT EpiLC (i.e. Group I enhancers) (Fig. 4b-c, Supplementary Fig. 4b). Moreover, the higher H3K4me1/2 levels observed in *Otx2^−/−^* EpiLC and EpiSC were not due to an increase in these histone modifications already in *Otx2^−/−^* ESC (Supplementary Fig. 4c). Nevertheless, the correlation between germline competence and H3K4me1 levels within PGCLC enhancers was not perfect, since *Otx2^−/−^* d4 EpiSC displayed higher germline competence than WT EpiLC, yet slightly lower H3K4me1 levels within Group I enhancers (Fig. 4c). Therefore, in addition to H3K4me1, other chromatin features within PGCLC enhancers might also contribute to the extended germline competence of *Otx2^−/−^* cells. In agreement with this possibility, Group I enhancers showed slightly higher H3K4me2 in *Otx2^−/−^* d4 EpiSC than in WT EpiLC (Fig. 4c). Furthermore, genome-wide as well as detailed analysis of representative enhancers (*i.e. Esrrb* and *Prdm1* enhancers) showed that the increased competence of *Otx2^−/−^* EpiLC and d4 EpiSC was also reflected in reduced CpG methylation levels within Group I PGCLC enhancers (Fig. 4d, Supplementary Fig. 4d). Overall, as the PGCLC genes get properly silenced in *Otx2^−/−^* EpiLC (Supplementary Fig. 1g), these results suggest that the extended germline competence of *Otx2^−/−^* cells could be linked to the impaired and delayed decommissioning of a subset of PGCLC enhancers.

**Fig. 4:**
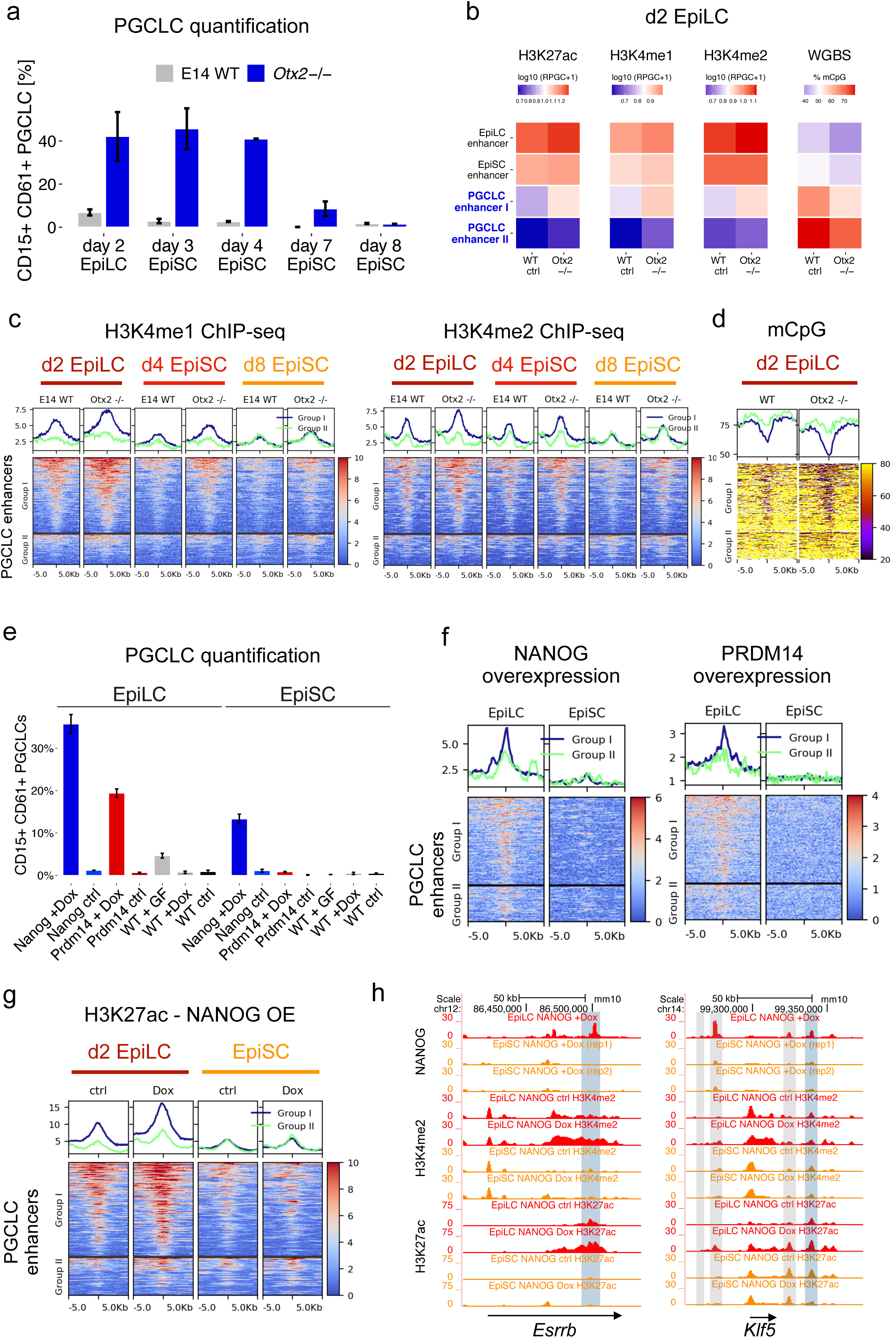
*In vitro* germline competence is correlated with the partial decommissioning and permissive chromatin state of PGCLC enhancers. a.) E14 WT and *Otx2*^−/−^ ESC were differentiated into d2 EpiLC, d4 EpiSC or d8 EpiSC, which were then further differentiated into PGCLC. PGCLC were quantified as the proportion of CD15^+^CD61^+^ cells found within d4 EB. The error bar represents the standard deviation from two biological replicates. b.) Summary plot showing the average levels of different epigenetic marks measured in WT and *Otx2*^−/−^ d2 EpiLC for the EpiLC, EpiSC and Group I+II PGCLC enhancers. Quantifications were performed by measuring the average signals of each epigenetic mark within −/+ 1kb of the enhancers. The H3K4me1 ChIP-seq data shown for WT d2 EpiLC is the same one used in Fig. 3c as second replicate. RPGC: Reads per genomic content. c.) Average profile (top) and heatmap (bottom) plots showing H3K4me1 and H3K4me2 levels for the Group I and Group II PGCLC enhancers in d2 EpiLC, d4 EpiSC and d8 EpiSC differentiated from either E14 WT or *Otx2*^−/−^ ESC. The H3K4me1 ChIP-seq data shown for WT d2 EpiLC and WT d8 EpiSC are the same ones used in Fig. 3c as second replicates. d.) Average profile (top) and heatmap (bottom) plots showing the percentage of CpG methylation for the Group I and Group II PGCLC enhancers in d2 EpiLC differentiated from either WT or *Otx2*^−/−^ ESC. e.) WT ESC and ESC lines with a Doxycycline-inducible system enabling the overexpression of exogenous NANOG (blue) or PRMD14 (red) were differentiated into d2 EpiLC and EpiSC. Next, d2 EpiLC and EpiSC were differentiated into PGCLC with (+Dox) or without (ctrl) Doxycycline and in the absence of growth factors. As a positive control, WT d2 EpiLC and EpiSC were differentiated with growth factors (WT+GF). The percentage of PGCLC (i.e. CD15^+^CD61^+^ cells) was measured in d4 EB from two biological replicates. f.) Exogeneous PRDM14 or NANOG tagged with HA were overexpressed for 18 hours in EpiLC and EpiSC. Then, ChIP-seq experiments were performed using an anti-HA antibody. The average profile (top) and heatmap (bottom) plots show the binding of the exogenous PRDM14 and NANOG within Group I and Group II PGCLC enhancers. g.) The ESC line enabling the inducible overexpression of exogenous NANOG-HA was differentiated into d2 EpiLC and EpiSC. Next, the cells were either left untreated (ctrl) or treated with Dox to overexpress NANOG as described in (f) and H3K27ac ChIP-seq experiments were performed. The average profile (top) and heatmap (bottom) plots show H3K27ac levels within Group I and Group II PGCLC enhancers. h.) Genome-browser view of the *Klf5* and *Esrrb* loci showing the binding of the exogenously induced NANOG-HA in both EpiLC and EpiSC as well as the H3K27ac and H3K4me2 signals in untreated (-Dox) and Dox-treated (i.e. NANOG-HA overexpression) d2 EpiLC and EpiSC. For the NANOG-HA ChIP-seq profiles in EpiSC the results of two independent biological replicates are shown to illustrate the reproducibility of the weak binding within the PGCLC enhancers. The previously deleted PGCLC enhancers (Fig. 2, Supplementary Fig. S2) are highlighted in blue and other PGCLC enhancers found within the same loci are indicated in gray.

### Partially decommissioned PGCLC enhancers are accessible and responsive to transcriptional activators in EpiLC

To investigate whether the partial decommissioning of PGCLC enhancers in EpiLC compared to EpiSC could have any functional consequences, we evaluated the accessibility of these enhancers to major PGCLC transcriptional activators. To this end, we generated clonal ESC lines in which HA-tagged PRDM14 or NANOG could be overexpressed upon addition of Doxycycline (Dox) (Supplementary Fig. 4e-f). In agreement with previous reports, the overexpression of either PRDM14 or NANOG upon differentiation of EpiLC into PGCLC yielded a high percentage of PGCLC in the absence of growth factors (Fig. 4e, Supplementary Fig. 4g)^12, 54^. In contrast, the overexpression of PRDM14 or NANOG upon differentiation from EpiSC resulted in considerably less PGCLC (Fig. 4e, Supplementary Fig. 4g). To assess whether PGCLC enhancers were differentially accessible to these TFs in EpiLC and EpiSC, we performed ChIP-seq experiments after a short induction of HA-tagged PRDM14 or NANOG in both cell types. Importantly, PGCLC enhancers, especially those belonging to Group I, were considerably more bound by PRDM14-HA and NANOG-HA in EpiLC than in EpiSC (Fig. 4f). Furthermore, NANOG-HA binding to the PGCLC enhancers resulted in increased H3K27ac and H3K4me2 levels in EpiLC but not in EpiSC, particularly within Group I enhancers (Fig. 4g-h, Supplementary Fig. 4h). Altogether, these results indicate that PGCLC enhancers, especially those belonging to Group I, are both accessible and responsive to transcriptional activators in EpiLC but not in EpiSC.

### Increased decommissioning of PGCLC enhancers in the absence of H3K4me1

Our data suggest that a significant fraction of PGCLC enhancers display permissive chromatin features in EpiLC that might contribute to their increased germline competence compared to EpiSC. These permissive features could be attributed, at least partly, to the persistence of H3K4me1 in EpiLC, as this histone mark could (i) protect PGCLC enhancers from mCpG and H3K9me2/3^52, 55–59^ and/or (ii) facilitate the recruitment of chromatin remodelling complexes ^40^. To directly assess the importance of H3K4me1 for germline competence and PGCLC specification, we used mESC catalytically deficient for MLL3 and MLL4 (dCD and dCT ESC lines; Fig. 5a)^38^. Previous work showed that in dCD ESC, active enhancers show a moderate reduction in H3K27ac, while gene expression, eRNA levels and the binding of MLL3/4 and their associated complexes are not affected, indicating that H3K4me1 is largely dispensable for the maintenance of enhancer activity (Supplementary Fig. 5a)^38, 39^. Additional characterization of dCD and dCT cell lines revealed that, upon differentiation, the major reductions in H3K4me1 and H3K4me2 that PGCLC enhancers already displayed in ESC became further exacerbated in EpiLC and EpiSC (Fig. 5b-c, Supplementary Fig. 5b-d). Moreover, H3K27ac levels within PGCLC enhancers were also reduced in dCD/dCT cells compared to their WT counterparts, although such differences were not as pronounced as for H3K4me1 (Fig. 5b, Supplementary Fig. 5b,d). Similarly, other enhancer groups (i.e. EpiLC and EpiSC enhancers) also displayed strong H3K4me1/2 losses and milder H3K27ac reductions in dCD cells (Fig. 5c, Supplementary Fig. 5d,g). In contrast, the levels of previous active histone modifications were rather similar around the TSS of PGCLC genes, with H3K4me1 even showing a slight increase in dCD cells (Supplementary Fig. 5d). In agreement with the protective role of H3K4me1/2 against CpG methylation^52, 56^, mCpG levels within PGCLC enhancers were generally elevated in dCD ESC^60^, while in dCD EpiLC the increased methylation was evident among some Group I enhancers (Fig. 5d, Supplementary Fig. 5e-f). Next, to evaluate the potential functional consequences of the previous epigenetic changes, we generated RNA-seq data in WT and dCD ESC, EpiLC and EpiSC. In agreement with previous work in ESC indicating that H3K4me1 is dispensable for enhancer function^38, 39^, the genes associated with PGCLC, EpiLC and EpiSC enhancers showed rather minor expression differences between WT and dCD cells in the three investigated cell types (Fig. 5e, Supplementary Fig. 5h-i, Supplementary data 4).

**Fig. 5:**
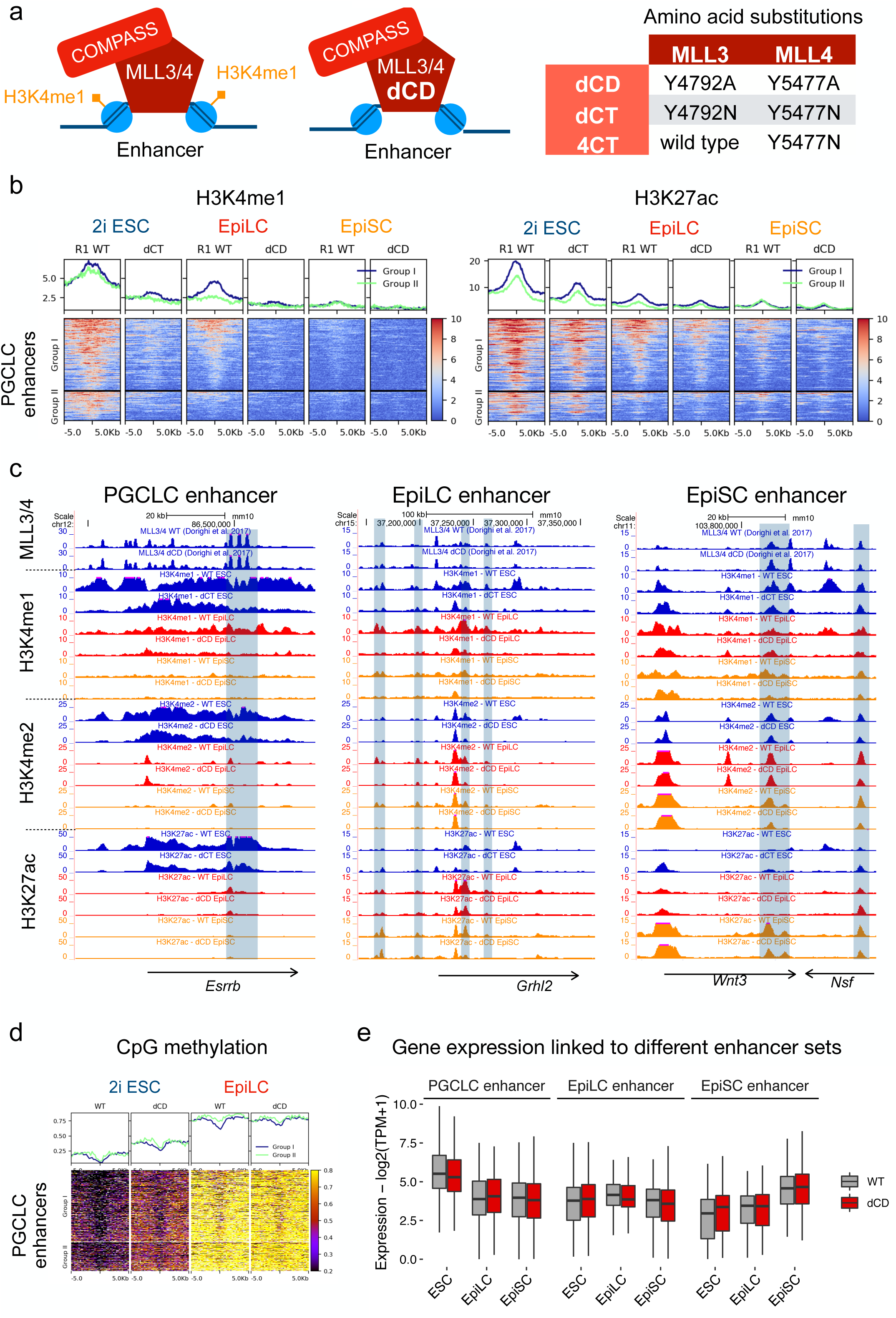
H3K4me1 is dispensable for enhancer-dependent gene expression in *in vitro* pluripotent cells: a.) Schematic illustration of the dCD cells, which express catalytic mutant MLL3 and MLL4 proteins without histone methyltransferase activity but are capable of binding to its target sites and interacting with other proteins as part of the COMPASS complex. The different *Mll3/4* catalytic mutants used in this study are shown to the right. b.) Average profile (top) and heatmap (bottom) plots showing H3K4me1 and H3K27ac levels for for the Group I and Group II PGCLC enhancers in R1 WT and MLL3/4 catalytic mutant ESC lines as well as upon their differentiation into d2 EpiLC and EpiSC. The H3K4me1 ChIP-seq data shown for WT d2 EpiLC and WT EpiSC are the same ones used in Fig. 3c as third replicates. c.) Genome-browser view showing H3K4me1, H3K4me2 and H3K27ac profiles in both WT and MLL3/4 catalytic mutant cells (ESC, EpiLC and EpiSC) around representative PGCLC (*Esrrb)*, EpiLC (*Grhl2*) and EpiSC (*Wnt3*) genes and their associated enhancers. d.) Average profile (top) and heatmap (bottom) plots showing the ratio of mCpG/CpG for the Group I and Group II PGCLC enhancers in WT and dCD ESC as well as upon their differentiation into d2 EpiLC. The WGBS data in 2i ESC was obtained from Skvortsova *et al.* 2019. e.) Box plots showing the expression, as measured by RNA-seq, of the genes associated with the PGCLC, EpiLC and EpiSC enhancers in WT (gray) and dCD (red) ESC cells as well as upon their differentiation into EpiLC and EpiSC. The RNA-seq data in ESC was obtained from Dorighi et al. 2017 and the experiments in d2 EpiLC and EpiSC were performed in duplicates.

Overall, our analyses indicate that the decommissioning of a subset of PGCLC enhancers gets exacerbated in dCD EpiLC compared to their WT counterparts, resulting in a chromatin state similar to the one observed in WT EpiSC (*i.e.* lower H3K4me1 and higher mCpG; Fig. 3). However, these epigenetic changes do not result in major gene expression changes in any of the investigated *in vitro* pluripotent cell types.

### H3K4me1 is necessary for *in vitro* germline competence

To address whether the increased decommissioning of PGCLC enhancers observed in dCD/dCT EpiLC could compromise their germline competence, these MLL3/4 catalytic mutant cell lines were differentiated into PGCLC. Chiefly, both dCD and dCT cells showed a significant reduction in their PGCLC differentiation capacity (Fig. 6a, Supplementary Fig. 6a). In agreement with MLL3 and MLL4 being functionally redundant^61^, such PGCLC differentiation defects were not observed when using cells that were catalytic mutant for MLL4 but not MLL3 (*i.e.* 4CT cells) (Fig. 5a, Fig. 6a). Furthermore, the compromised PGCLC differentiation of the dCD cells was still observed at a later time point (day 6, Supplementary Fig. 6b), indicating that the observed defects are not simply explained by a delay in PGCLC specification.

**Fig. 6:**
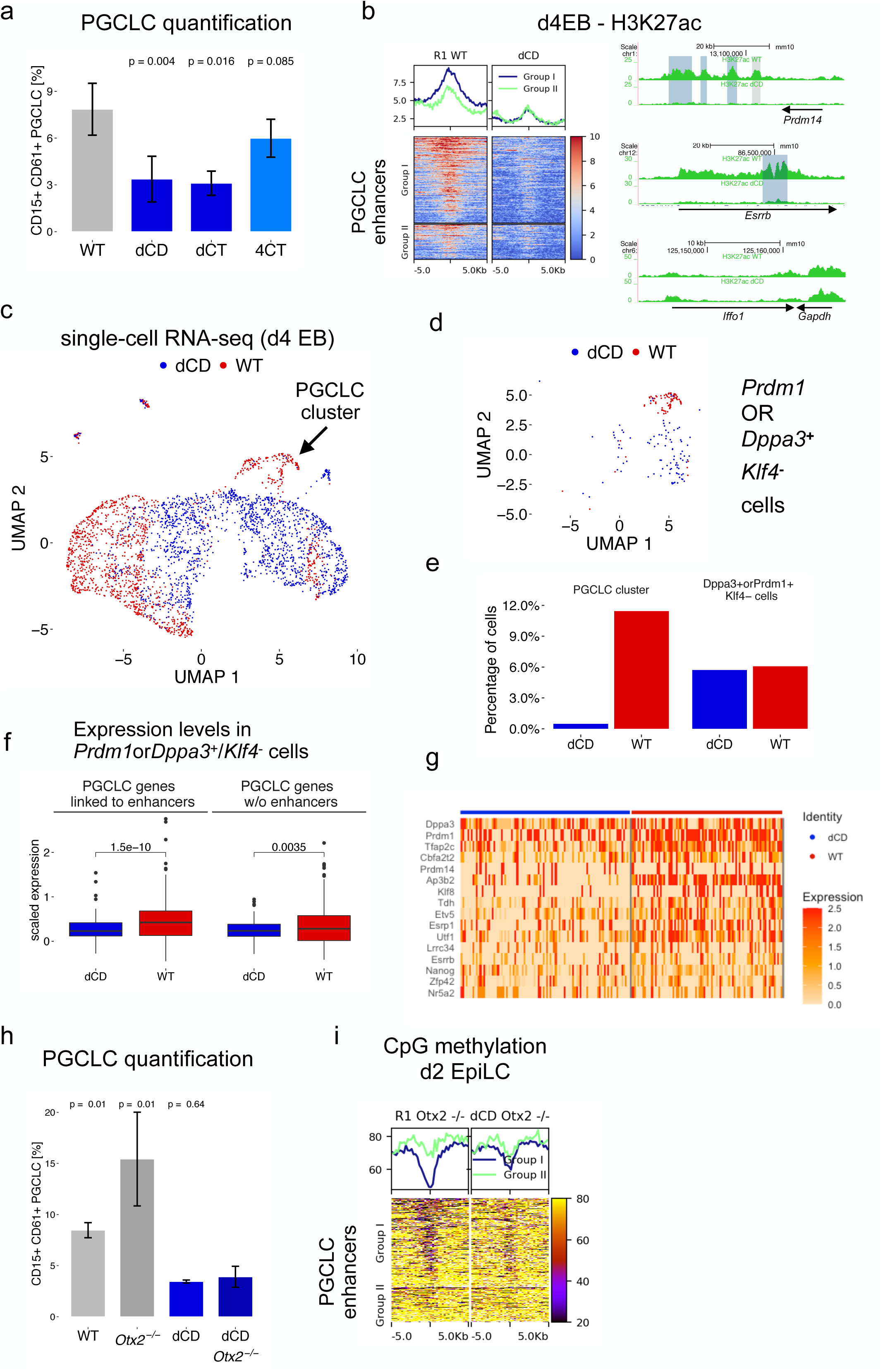
H3K4me1 is required for the proper induction of PGCLC and their associated gene expression program. a.) WT ESC and ESC lines with the indicated MLL3/4 amino acid substitutions were differentiated into PGCLC. PGCLC were defined as CD15^+^CD61^+^ cells within d4 EB. Each PGCLC quantification was performed in at least four biological replicates. The p-values were calculated using Wilcoxon tests. b.) H3K27ac ChIPmentation experiments were performed in d4 EB differentiated from R1 WT and dCD ESC. Left: Average profile (top) and heatmap (bottom) plots showing H3K27ac levels for the Group I and Group II PGCLC enhancers. Right: Genome browser views showing H3K27ac profiles in R1 WT and dCD d4 EB around two representative PGCLC genes and their linked enhancers (*Esrrb* and *Prdm14*). The CRISPR-Cas9 deleted enhancer regions (Fig. 2) are highlighted in blue and other PGCLC enhancers found within the same loci are indicated in gray. Lastly, H3K27ac profiles are shown around a housekeeping gene (*Gapdh*) to illustrate the comparable quality of the H3K27ac ChIP-seq data generated in both WT and dCD d4 EB. c.) UMAP plot of the single-cell RNA-sequencing data generated in d4 EB differentiated from R1 WT and dCD ESC. The PGCLC cluster, which was identified by the expression of PGCLC markers is indicated with an arrow. Each red and blue dot in the plot represents a WT and dCD individual cell, respectively. d.) Cells expressing *Prdm1* or *Dppa3* but not the naïve pluripotency marker *Klf4* (*Prdm1orDppa3+/Klf4-*) in WT (red) and dCD (blue) d4 EBs are displayed within the same UMAP plot shown in (c). e.) Quantification of the percentage of WT and dCD d4 EB cells found within the PGCLC cluster (Fig. 6c and Supplementary Fig. 6c) or expressing *Prdm1/Dppa3* but not *Klf4* (*Prdm1orDppa3+/Klf4-)* (Fig. 6d). f.) Box plots showing the expression of the PGCLC genes linked (left, n=216) or not (right, n=170) to at least one PGCLC enhancer within WT (red) and dCD (blue) *Prdm1orDppa3+/Klf4-* cells. WT and dCD were compared using a paired student t-test. g.) Heatmap showing the expression of selected PGCLC genes (rows) linked to PGCLC enhancers within individual WT and dCD *Prdm1orDppa3+/Klf4-* cells (columns). h.) R1 WT, *Otx2*^−/−^, dCD and dCD *Otx2*^−/−^ ESC were differentiated into PGCLC. PGCLC were quantified as the proportion of CD15^+^CD61^+^ cells found within d4 EB. The PGCLC differentiations for the dCD *Otx2*^−/−^ ESC were performed in three biological replicates and using two different clonal lines (n=3×2). The other PGCLC measurements were performed in biological triplicates. i.) Whole-genome bisulfite sequencing experiments were performed in d2 EpiLC differentiated from R1 *Otx2*^−/−^ and dCD *Otx2*^−/−^ ESC. The average profile (top) and heatmap (bottom) plots show the percentage of CpG methylation for the Group I and Group II PGCLC enhancers in WT and dCD d2 EpiLC.

Next, we investigated whether the reduced germline competence of the dCD cells could be caused by compromised PGCLC enhancer reactivation upon PGCLC differentiation. H3K27ac ChIP-seq profiling of d4 EB derived from WT and dCD ESC (Fig. 6b) showed that PGCLC enhancers, especially those belonging to Group I, displayed higher H3K27ac levels in WT than in dCD d4 EB. Since the previous ChIP-seq experiments were performed in EBs and not in sorted PGCLC, the low H3K27ac levels in dCD cells could be caused by either a defect in the activation of PGCLC enhancers and their associated genes or by an overall reduction in the number of PGCLC present within d4 EBs. To distinguish between these two possibilities, we performed scRNA-seq analyses of WT (1416 cells) and dCD (1699 cells) d4 EBs (Supplementary Data 5). UMAP (Uniform Manifold Approximation and Projection for Dimension Reduction) analysis of the resulting single cell transcriptomes confirmed the presence within d4 EB of subclusters resembling the main cell populations (*i.e.* PGCLC, ExEctorderm-like, ExMesoderm-like) that characterize the proximo-posterior end of the mouse embryo following germline specification (Supplementary Fig. 6c-d). Most importantly, the PGCLC cluster consisted mostly of WT cells (Fig. 6c,e), thus in agreement with the results obtained using FACS and cell surface markers (Fig. 6a). We also identified subclusters (*i.e.* ExEndoderm/Gut-like, 2-cell-like) that were not observed in previous scRNA-seq analyses (Fig. 1, Supplementary Fig. 1), probably due to the lower number of d4 EB cells sequenced in those initial experiments. Moreover, while the transcriptomes of some extraembryonic-like cell types were similar between WT and dCD cells (e.g. Ex-Endoderm/Gut-like, Endothelial-like), many dCD cells were located in clusters with poorly defined identity that did not show differential expression of specific markers. These “undefined” clusters were transcriptionally more heterogeneous than the remaining clusters found within d4 EB (Supplementary Fig. 6e). Interestingly, we noticed that within the dCD EBs there were cells expressing major PGC markers (i.e. *Prdm1* or *Dppa3*) but not naïve pluripotency ones (i.e. *Klf4*) (Fig. 6d), suggesting a cellular identity similar to PGCLC. The proportion of these *Prdm1orDppa3+/Klf4-* cells was similar among WT and dCD EBs (Fig. 6e). However, while in the WT EBs these cells were mostly found within the PGCLC subcluster, in the dCD EBs they were part of the subclusters with poorly defined identity (Fig. 6e), suggesting important transcriptional differences between WT and dCD *Prdm1orDppa3+/Klf4-* cells. Congruently, the expression of the PGCLC genes associated with PGCLC enhancers (*e.g. Tfap2c*, *Prdm14*, *Utf1*, *Esrrb*) was significantly reduced in dCD *Prdm1orDppa3+/Klf4-* cells in comparison to their WT counterparts, while no significant differences were observed for the PGCLC genes without associated enhancers (Fig. 6f-g). Overall, these scRNA-seq analyses suggest that the induction of the PGCLC expression program, particularly of those genes linked to PGCLC enhancers, is compromised in dCD cells.

Finally, to further assess the importance of H3K4me1 for PGCLC induction, we investigated whether the extended germline competence of *Otx2^−/−^* EpiLC could be also attributed, at least partly, to the retention of H3K4me1 and the impaired decommissioning of PGCLC enhancers (Fig. 4). To this end, we deleted *Otx2* in dCD ESC (*i.e.* dCD *Otx2^−/−^*) as well as in their parental WT ESC (*i.e.* R1 *Otx2^−/−^*) (Supplementary Fig. 6f) and differentiated them into PGCLC. As expected, the deletion of *Otx2* in the R1 ESC resulted in increased germline competence (Fig. 6h), although not as pronounced as in E14 ESC (Fig. 4a)^20^, which could be attributed to the variable germline competence observed among different ESC lines^10^. Most importantly, both dCD and dCD *Otx2^−/−^* EpiLC showed a strong and similar reduction in their PGCLC differentiation capacity (Fig. 6h). Furthermore, genome-wide and locus-specific analyses of mCpG levels revealed that Group I PGCLC enhancers were considerably more methylated in dCD *Otx2^−/−^* EpiLC than in *Otx2^−/−^* EpiLC (Fig. 6i, Supplementary Fig. 6g).

Altogether, our data shows that H3K4me1 is required for *in vitro* germline competence and proper PGCLC induction. Although we cannot rule out that gene expression and epigenetic changes in ESC and/or extraembryonic-like cell types ^62^ might also contribute to the PGCLC differentiation defects observed in dCD/dCT cells, our data suggests that the persistence of H3K4me1 within the PGCLC enhancers might facilitate their reactivation during PGCLC induction.

### PGCLC enhancers get heterogeneously decommissioned in formative epiblast cells *in vivo*

Overall, our work using the PGCLC *in vitro* differentiation system suggests that germline competence entails the persistence of permissive chromatin features within a significant fraction of PGCLC enhancers. To evaluate whether this is also true *in vivo*, we took advantage of several epigenomic datasets recently generated from mouse embryos^63^. Firstly, analysis of DNAse I data obtained in E9.5 and E10.5 PGC showed that our *in vitro* defined PGCLC enhancers are highly accessible *in vivo* (Fig. 7a). Next, to evaluate the dynamics of PGCLC enhancer decommissioning, we analyzed genome-wide CpG methylation data from mouse epiblasts^64^. In agreement with our *in vitro* observations, PGCLC enhancers showing incomplete decommissioning in EpiLC (i.e. Group I enhancers) displayed lower CpG methylation levels in germline competent E5.5 epiblast cells than in the E6.5 epiblast (Fig. 7b), in which germline competence is already reduced^4^.

**Fig. 7:**
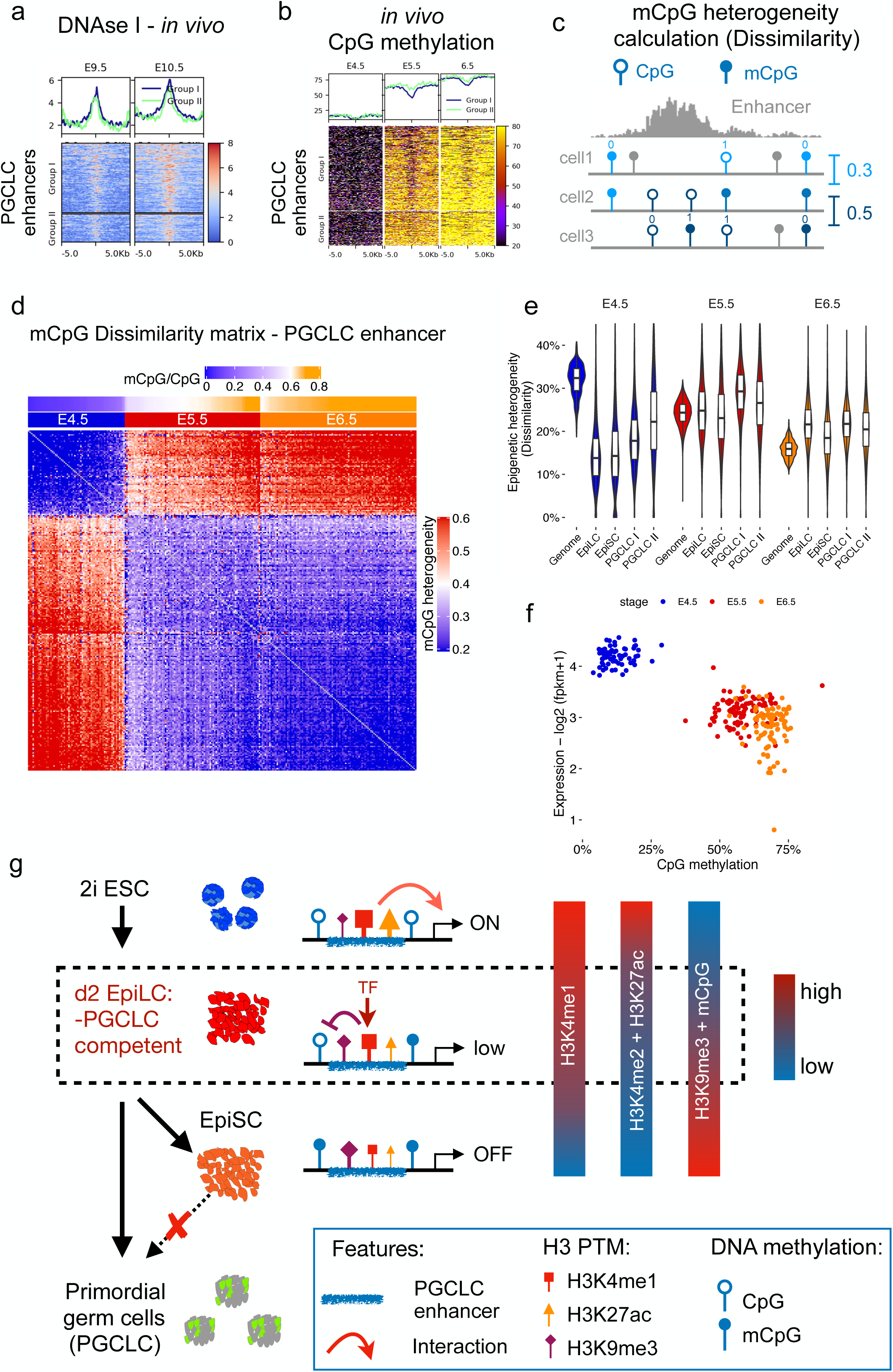
Chromatin features and epigenetic heterogeneity of PGCLC enhancers *in vivo.* a.) Average profile (top) and heatmap (bottom) plots showing DNAse-seq levels for the Group I and Group II PGCLC enhancers in PGCs isolated from E9.5 and E10.5 mouse embryos. The DNAse-seq data were obtained from Li *et al.* 2018. b.) Average profile (top) and heatmap (bottom) plots showing the percentage of CpG methylation for the Group I and Group II PGCLC enhancers in E4.5, E5.5 and E6.5 mouse epiblasts. The genome-wide mCpG data were obtained from Zhang *et al.* 2018. c.) Illustration of the mCpG heterogeneity estimation, which is based on the mCpG dissimilarity concept. Briefly, for each pairwise comparison, the methylation status of only those CpGs that are covered in the two cells being compared is considered (blue lollipops). Then, if two cells show the same methylation status they receive a value of 0 (similarity) and if they show dissimilar methylation patterns a value of 1. The mean of all pairwise comparisons reflect the dissimilarity of the CpG methylation, or mCpG heterogeneity. d.) CpG methylation heterogeneity heatmap showing the differences in mCpG within PGCLC enhancers between pairs of individual cells. The CpG methylation heterogeneity values are presented with a blue-red scale (blue means that cells are similar in their methylation status, while red indicates that they are more dissimilar/ heterogenous). Above the heatmap, the developmental stages of the investigated cells (E4.5, E5.5 or E6.5) and the average CpG methylation (blue-orange scale) measured for all PGCLC enhancers within each single cell are shown (n=261 cells). For each developmental stage, the cells were ranked according to the average mCpG levels within the PGCLC enhancers. e.) Violin plots showing the CpG methylation heterogeneity measured in E4.5, E5.5 and E6.5 epiblast cells for all covered CpGs between two cells in the mouse genome (genome-wide) as well as within EpiLC, EpiSC and PGCLC enhancers (Group I and II). e.) Comparison of the single-cell RNA expression and CpG methylation of E4.5, E5.5 and E6.5 epiblasts. Plotted is the average expression of genes linked to Group I PGCLC enhancers and the average CpG methylation of these enhancers (n=256). f.) Model illustrating the partial decommissioning of Group I PGCLC enhancers during PGCLC differentiation and its relevance for germline competence. Most of the Group I PGCLC enhancers are already active in ESC (i.e. high H3K27ac and H3K4me1/2 levels). Upon differentiation into EpiLC and EpiSC, these enhancers progressively lose these active histone modifications, albeit with different temporal dynamics: (i) H3K27ac and H3K4me2 are rapidly lost and are already low germline competent EpiLC; (ii) H3K4me1 partially persist in EpiLC and is significantly lower in EpiSC. On the other hand, repressive epigenetic features, such as H3K9me3 and CpG methylation, start accumulating within the PGCLC enhancers already in EpiLC, but they increase even further in EpiSC. Using H3K4me1 deficient cells, we show that the persistence of H3K4me1 within PGCLC enhancers facilitates their reactivation during PGCLC induction, thus suggesting that this histone modification can be important for *in vitro* germline competence. Mechanistically, the persistence of H3K4me1 within PGCLC enhancers might protect them from CpG methylation and render them accessible and responsive to transcriptional activators.

Previous single-cell CpG methylation analyses revealed that, during mouse peri-implantation stages, the formative epiblast is particularly heterogeneous, especially within enhancers^65^. Therefore, we hypothesize that the lower CpG methylation levels observed for some PGCLC enhancers in the E5.5 epiblast in comparison to the E6.5 epiblast could be the result of increased cell-to-cell variation. To evaluate this idea, we analyzed single-cell data in which DNA methylation and gene expression were measured for the same cells across different epiblast stages (*i.e.* E4.5, E5.5 and E6.5)^43^. Firstly, we measured mCpG heterogeneity by comparing the methylation status of individual CpGs within PGCLC enhancers^66^ (Fig. 7c). This analysis revealed that the formative epiblast (E5.5) displayed the highest variation in mCpG, while in the primed epiblast (E6.5) the PGCLC enhancers were more homogeneously methylated (Fig. 7d). Interestingly, when comparing different enhancer sets across epiblast stages, the highest epigenetic heterogeneity (∼30 %) was observed for the Group I PGCLC enhancers in the E5.5 epiblast (Fig. 7e). As the CpG coverage for the Group I PGCLC enhancers was similar across epiblast stages, the previous differences in mCpG heterogeneity are unlikely to be caused by technical reasons (Supplementary Fig. 7a). Lastly, we compared mCpG levels within Group I PGCLC enhancers and the expression of their associated genes across the profiled epiblast cells (Fig. 7f). Similar to what we observed for EpiLC and EpiSC, the PGCLC genes associated with the Group I enhancers display low and similar expression in E5.5 and E6.5 epiblast cells despite the differences in CpG methylation.

Overall, the previous data support that PGCLC enhancers are also partially decommissioned in the formative epiblast and indicate that such partial decommissioning could indeed reflect epigenetic heterogeneity. Nevertheless, the relevance of the partial decommissioning and epigenetic heterogeneity of PGCLC enhancers for germline competence *in vivo* remains to be demonstrated.

## DISCUSSION

Direct evidence supporting the functional relevance of H3K4me1 are scarce, partly due to the difficulties in separating the enzymatic and non-enzymatic functions of histone methyltransferases^67^. This limitation was recently overcome by establishing *Mll3/4* catalytic mutant ESC lines in which H3K4me1 is lost from enhancers^38^. Under self-renewing conditions, the loss of H3K4me1 partly reduced H3K27ac but did not affect transcription from either enhancers or gene promoters, suggesting that H3K4me1 is dispensable for the maintenance of enhancer activity^38^. Similarly, H3K4me1 was not required for *de novo* enhancer activation (*i.e.* without prior presence of H3K4me1) upon somatic differentiation of ESC (Fig. 5)^62^. However, it is still unclear whether H3K4me1 is important for the activation of primed enhancers already pre-marked with this histone modification^25, 31^. In the case of *in vitro* germline competence, here we report that a subset of PGCLC enhancers gets partly decommissioned in EpiLC and retains permissive chromatin features, including H3K4me1, already present in a preceding active state (*i.e.* in 2i ESC) (Fig. 7g). This resembles the so-called latent enhancers previously described in differentiated macrophages, in which, following an initial round of activation and silencing, the persistence of H3K4me1 was proposed to facilitate subsequent enhancer induction upon restimulation^27^. The mechanisms involved in the persistence of H3K4me1 and other permissive chromatin features are still unknown, although we can envision at least two non-mutually exclusive possibilities: (i) a passive mechanism whereby MLL3/4 binding to PGCLC enhancers is already lost in EpiLC, but H3K4me1 can still be transiently retained due to the slow dynamics of H3K4 demethylation^68, 69^; (ii) an active maintenance mechanism similar to the one reported for enhancer priming^24, 25^, whereby the binding of certain TFs might enable the persistent recruitment of MLL3/4 and the retention of H3K4me1 within PGCLC enhancers. Since PGCLC enhancers display low and similar ATAC-seq signals in EpiLC and EpiSC (Fig. 3), this would argue in favor of passive mechanisms rather than an active retention of TFs and co-activators (e.g. MLL3/4) in EpiLC.

*Drosophila melanogaster* embryos expressing catalytic deficient *Trr*, the homolog of mammalian *Mll3/4,* develop normally, thus arguing against a major role for H3K4me1 in enhancer function^39^. In contrast, the loss of MLL3/4 catalytic activities in mouse embryos resulted in early embryonic lethality (∼E8.5)^62^, suggesting that H3K4me1 is functionally relevant during mammalian embryogenesis. It is important to consider that, in comparison to mammalian cells, the *D. melanogaster* genome is largely devoid (< 0.03 %) of CpG methylation throughout its entire life cycle^70^. Previous studies have extensively documented the protective role of H3K4me1/2 against mCpG and heterochromatinization^52, 55–59^. Congruently, we showed that H3K4me1 and mCpG levels within PGCLC enhancers are negatively correlated and that H3K4me1-deficient cells displayed increased mCpG levels within Group I PGCLC enhancers. Therefore, although the importance of mCpG for enhancer activity is still debated^71–73^, it is tempting to speculate that the lack of DNA methylation in flies render them less sensitive to H3K4me1 loss. Nevertheless, *Trr* catalytic mutant fly embryos displayed aberrant phenotypes under stress conditions, suggesting that H3K4me1 might help to fine-tune enhancer activity under suboptimal conditions. Similarly, we found that the induction of PGCLC genes linked to enhancers was disrupted in dCD cells. Therefore, we propose that H3K4me1 might facilitate, rather than being essential for, enhancer (re)activation and the robust induction of developmental gene expression programs (Fig. 7g). This H3K4me1 facilitator role might involve not only protection from DNA methylation and heterochromatinization but also increased accessibility and responsiveness to transcriptional activators (Fig. 4)^40^ as well as physical proximity between genes and enhancers^32^. Future work in additional species and cellular transitions will establish the relevance and prevalence of H3K4me1 function within enhancers.

Finally, our work highlights the diverse mechanisms whereby the epigenetic status of enhancers can contribute to developmental competence. For instance, a subset of early brain enhancers displays a poised state in ESC (*i.e*. H3K4me1+H3K27me3) that might facilitate their activation upon neural differentiation^74^, while in endodermal progenitors the binding of pioneer TFs and the consequent priming of relevant enhancers by H3K4me1 might facilitate their activation during subsequent differentiation^24, 25^. Moreover, recent work based on chromatin accessibility or multi-omic profiling indicates that the commitment towards certain cell lineages involves the activation of already accessible enhancers, while in other cases this can occur through *de novo* enhancer activation^43, 75^. Future work will elucidate the prevalence and regulatory mechanisms by which the epigenetic state of enhancers can contribute to cellular competence and the robust deployment of developmental gene expression programs. In this regard, it would be important to evaluate whether, as reported here for PGCLC induction, the partial decommissioning of enhancers can be involved in their subsequent reactivation and, thus, in the induction of gene expression programs in other developmental contexts. Similar mechanisms might be also important in other physiological^27^ and pathological^76, 77^ contexts in which a previously used but already dismantled gene expression program gets re-activated.

## ACKNOWLEDGEMENTS

We thank the Rada-Iglesias lab members for insightful comments and critical reading of the manuscript, Antonio Simeone and Christa Buecker for generously providing the *Otx2^−/−^* cell line.

Library preparations and next-generation sequencing experiments were performed in the NGS Core Facility of the Cologne Center for Genomics (CCG). Computational analyses were performed on the Cologne High Efficient Operating Platform for Science (CHEOPS).

Cell sorting and Flow cytometry experiments were performed in the FACS Facility of the Centre for Molecular Medicine (CMMC) and the FACS & Imaging Core Facility at the Max Planck Institute for Biology of Ageing.

## Author Contributions

Conceptualization, T.B., A.R-I.; Experimental investigation, T.B., K.S., G.C., P.R., M.B., S.J.C, L.B., A.B., M.L, K.D; Data analysis: T.B., G.C.; Writing, Review & Editing, T.B., A.R-I; Resources, W.F.J. vI, M.M., J.W., W.R., A.R.-I.; Supervision and Funding Acquisition, A.R.-I.;

## Methods

### Data availability

The sequencing datasets generated during this study are available through GEO (GSE155089): https://www.ncbi.nlm.nih.gov/geo/query/acc.cgi?acc=GSE155089

Codes used in the computational analyses of the sequencing data are available through *GitHub*: https://github.com/ToreBle/Germline_competence.

UCSC sessions to visualize all the processed sequencing data are available using the links provided in the “*ChIP-seq data processing”* section (see below).

### WT and transgenic ESC lines

Two different male mouse ESC lines (*i.e.* E14Tg2a and R1) were used as indicated for each particular experiment. The *Mll3/4* dCD, *Mll3/4* dCT and *Mll4* CT ESC lines were derived from WT R1 ESC as described in^38^. The E14 *Otx2^−/−^* ESC line^53, 78^ and its parental E14Tg2a ESC were kindly provided by Christa Buecker. Other transgenic ESC lines generated in this study are more extensively described in following sections and are derived from E14Tg2a ESC.

### Cell culture and differentiation protocols

All ESC lines were cultured under serum+LIF conditions both for regular maintenance and CRISPR/Cas9 genome editing experiments. More specifically, ESC were cultured on gelatin-coated plates using Knock-out DMEM (*Life Technologies*) supplemented with 15% FBS (*Life Technologies*) and LIF. Before each PGCLC differentiation, ESC were adapted to 2i media (serum-free N2B27 medium supplemented with MEK inhibitor PD0325901 [0.4 μM, *Miltenyi Biotec*], GSK3β inhibitor CHIR99021 [3 μM, *Amsbio*], and LIF) for at least four days in gelatin-coated tissue culture plates^79^. All measurements described in the manuscript for ESC were performed under 2i conditions.

For EpiLC and PGCLC differentiation, the protocols from^79^ were followed. For the EpiLC differentiation, 6-well plates were coated with 15 µg ml^−1^ Fibronectin and 2 × 10^5^ 2i ESC were differentiated in N2B27 media supplemented with 20 ng ml^−1^ Activin A, 12 ng ml^−1^ bFGF and 10 µl ml^−1^ KSR for two days (unless indicated otherwise for particular experiments). For the PGCLC differentiation, EpiLC or EpiSC were plated at a density of 2 × 10^4^ cells ml^−1^ and cultured as embryoid bodies (EB) on Ultra-Low attachment multiwell plates (*Corning® Costar®*) in GK15 medium supplemented with growth factors (0.5 µg ml^−1^ BMP4, 0.1 µg ml^−1^ SCF, 1000 U ml^−1^ LIF and 50 ng ml^−1^ EGF, no BMP8a). Unless otherwise indicated, PGCLC were typically analyzed after four days.

EpiSC differentiation was performed according to^80^. 2i ESC were plated at a density of 2.5 × 10^5^ cells ml^−1^ on plates coated with 15 µg ml^−1^ Fibronectin and differentiated into EpiSC by passaging them in N2B27 media supplemented with 20 ng ml^−1^ Activin A and 12 ng ml^−1^ bFGF for at least eight days or as indicated in the results section for particular experiments.

### Enhancer and gene deletions with CRISPR/Cas9

Pairs of gRNAs flanking each of the selected enhancers/genes were designed with *benchling* (https://www.benchling.com). ESC were transfected with pairs of gRNA-Cas9 expressing vectors^81^ specific for each enhancer/gene using Lipofectamine 3000 (*invitrogen*). After transfection and puromycin selection, single cells were seeded into a 96 well plate to derive clonal cell lines. Then, clonal lines were genotyped by PCR and the presence of the intended deletions in each clonal line was verified by Sanger-sequencing. gRNA sequences and genomic coordinates of the different deletions are listed in Supplementary Data 6.

### Generation of the Mll3/4 dCD, Mll3/4 dCT and Mll4 CT ESC lines

The *Mll3/4* dCD ESC line was previously generated and described^38^. The *Mll3/4* dCT and *Mll4* CT ESC lines were also generated as described in^38^. Briefly, R1 ESC were co-transfected with a 200 bp single stranded oligonucleotide donor template harboring the desired point mutations and a vector coexpressing Cas9 and gRNAs targeting *Mll3* and *Mll4*. After 48 hr, single GFP+ cells were sorted into 96 well plates coated with fibronectin (5 mg ml^−1^). The resulting clonal ESC lines were genotyped by PCR amplification of a region spanning the cleavage site, followed by digestion with a restriction enzyme predicted to only cut the wild-type PCR products. Once candidate ESC lines were selected, the presence of the intended *Mll3/4* mutations was confirmed by Sanger sequencing.

### Ectopic and inducible expression of candidate transcription factors

Mouse *Prdm14* and *Nanog* cDNAs were amplified and cloned into a Doxycycline (Dox)-inducible piggyBac vector^82^ enabling the ectopic expression of selected genes fused with a HA-tag at their C-terminus. The resulting piggyBac vectors were transfected together with a Super PiggyBac Transposase expressing vector (*Systems Bioscience*) and a Tet transactivator. After transfection, single cells were seeded into a 96 well plate to derive clonal ESC lines. The clonal ESC lines with the lowest expression of the transgenic genes in the absence of Dox were selected. To evaluate the effects of PRDM14 and NANOG ectopic expression on PGCLC specification, 1 µg ml^−1^ Dox was added once PGCLC differentiation was started. To investigate the effects of the ectopic expression of PRDM14 or NANOG in EpiLC and EpiSC, 1 µg ml^−1^ Dox was added 18 hours before the cells were collected for downstream ChIP-seq experiments (*i.e.* EpiLC differentiation was initiated, after 30 hours Dox was added and cells were collected after 48 hours; EpiSC were maintained in differentiation media for at least eight passages and then treated with Dox for 18 hours).

### Quantification of PGCLC by flow cytometry

In general, PGCLC were quantified using antibody staining and flow cytometry^79^. Briefly, after four days of PGCLC differentiation, the resulting EB were dissociated and stained for 45 minutes with antibodies against CD61 (*Biolegends*) and CD15 (*eBioscience*) conjugated with PE and Alexa Fluor 647, respectively.

All PGCLC quantifications were performed using the FACS CantoII Cytometer (*BD Bioscience*) equipped with the BD FACSDiva Software. PGCLC sorting was performed on a FACS AriaIII cell sorter (*BD Bioscience*).

### ATAC-seq

The ATAC-seq protocol was adapted from^83^ using 3.8 × 10^4^ and 5.0 × 10^4^ cells for the two replicates generated for each of the investigated cell types, respectively. Briefly, the cells were lysed in lysis buffer (10 mM Tris-HCl, pH 7.4, 10 mM NaCl, 3 mM MgCl_2_, 0.1% IGEPAL CA-630) supplemented with freshly added protease inhibitor for 15 min. Following centrifugation the pellet was resuspended in a Tn5 transposase reaction mix (*Illumina*) for 30 min at 37°C. Following DNA purification with the MinElute PCR purification kit (*Qiagen*), libraries were prepared with the Nextera DNA library prep kit (*Illumina*).

### ChIP-seq and ChIPmentation

Cells were cross-linked with 1 % formaldehyde for 10 min, followed by quenching with 0.125 M glycine, harvesting and washing in PBS containing protease inhibitors. The cells were sequentially lysed in three buffers (lysis buffer 1: 50 mM HEPES, 140 mM NaCl, 1 mM EDTA, 10% glycerol, 0.5% NP-40, 0.25% TX-100; lysis buffer 2: 10 mM Tris, 200 mM NaCl, 1 mM EDTA, 0.5 mM EGTA; lysis Buffer 3: 10 mM Tris, 100 mM NaCl, 1 mM EDTA, 0.5 mM EGTA, 0.1% Na-Deoxycholate, 0.5% N-lauroylsarcosine) with rotation for 10 min in between. Then, the chromatin was sonicated with the Epishear™ Probe Sonicator (*Active Motif*) with 20 s ON and 30 s OFF for 8 cycles. After centrifugation, the supernatant was divided into input and ChIP samples. The ChIP samples were incubated with specific antibodies (Supplementary Data 6) overnight, followed by immunoprecipitation with Dynabeads Protein G beads (*invitrogen*). Next, the beads were washed with RIPA buffer (50 mM Hepes, 500 mM LiCl, 1 mM EDTA, 1% NP-40, 0.7% Na-Deoxycholate) on a magnet, eluted (50 mM Tris, 10 mM EDTA, 1% SDS), and reverse crosslinked at 65 °C overnight in parallel with the input. Finally, DNAs were purified with the ChIP DNA Clean & Concentrator (*Zymo Research*) and ChIP libraries were generated with the TruSeq kit (*Illumina*).

When ChIP-seq profiles were generated from low cell numbers (e.g. d4 EB obtained with the PGCLC differentiation protocol), the ChIPmentation protocol described by Schmidl *et al*. 2015^84^ was used. Following immunoprecipitation with Dynabeads Protein G beads (*invitrogen*) in PCR tubes, samples were subject to Tagmentation (5 µl Tagmentation Buffer, 1 µl Tagmentation DNA Enzyme (*Illumina*), 19 µl Nuclease free water) in order to incorporate sequencing adapters. Lastly, DNAs were eluted from the beads and used for library preparation with the Nextera DNA library preparation kit (*Illumina*).

### Locus-specific bisulfite sequencing

Genomic DNA was purified with phenol-chloroform and subjected to bisulfite conversion according to the EZ DNA Methylation-Direct™ Kit (Zymo Research). Then, primer pairs specific for each investigated enhancer were used for PCR amplification with the EpiTaq HS polymerase (*TaKaRa*). Finally, the resulting amplicons were gel-purified, subjected to blunt end cloning and analyzed by Sanger sequencing. The target sequence and the bisulfite primer for the selected PGCLC enhancers are listed in the Supplementary Data 6.

### ChIP-bisulfite sequencing

ChIP-bisulfite experiments were performed as described in^85^ with slight modifications. Firstly, the ChIP protocol described above was followed. After the final DNA purification, all the resulting H3K4me1 and H3K4me2 ChIP DNAs and 200 ng of the corresponding input DNA were subjected to bisulfite conversion according to the EZ DNA Methylation-Direct™ Kit (*Zymo Research*). Then, the subsequent amplification, purification and sequencing steps were performed as described for the locus-specific bisulfite sequencing experiments.

### Genome-wide bisulfite sequencing

Bisulfite sequencing libraries were prepared from column-purified DNA of d2 EpiLC using the PBAT method previously described in Clark *et al*. 2017^86^. Briefly, for the bisulfite conversion the instructions of the EZ Methylation Direct MagPrep Kit (*Zymo*) were followed. After purification, bisulfite converted DNAs were eluted from MagBeads directly into 39 µl of first strand synthesis reaction mastermix (1x Blue Buffer (*Enzymatics*), 0.4 mM dNTP mix (*Roche*), 0.4 µM 6NF preamp oligo (*IDT*)), heated to 65 °C for 3 minutes and cooled on ice. 50 U of klenow exo-(*Enzymatics*) was added and the mixture incubated on a thermocycler at 37 °C for 30 minutes after slowly ramping from 4 °C. Reactions were diluted to 100 µl and 20 U of exonuclease I (*NEB*) added and incubated at 37 °C before purification using a 0.8:1 ratio of AMPure XP beads. Purified products were resuspended in 50 µl of second strand mastermix (1x Blue Buffer (*Enzymatics*), 0.4 mM dNTP mix (*Roche*), 0.4 µM 6NR adaptor 2 oligo (*IDT*) then heated to 98 °C for 2 minutes and cooled on ice. 50 U of klenow exo-(*Enzymatics*) was added and the mixture incubated on a thermocycler at 37 °C for 90 minutes after slowly ramping from 4 °C. Second strand products were purified using a 0.8:1 ratio of AMPure XP beads and resuspended in 50 µl of PCR master mix (1x KAPA HiFi Readymix, 0.2 µM PE1.0 primer, 0.2 µM iTAG index primer) and amplified with 12 cycles. The final libraries were purified using a 0.8:1 volumetric ratio of AMPure XP beads before pooling and sequencing. All libraries were prepared in parallel with the pre-PCR purification steps carried out using a Bravo Workstation pipetting robot (*Agilent Technologies*).

### RNA isolation, cDNA synthesis and RT-qPCR

Total RNA from ESC, EpiLC and EpiSC was extracted following the protocol of the innuPREP DNA/RNA mini kit (*Analytik Jena*), while for the RNA extraction from d2 EB, d4 EB and sorted PGCLC the ReliaPrep™ RNA Miniprep Systems (*Promega*) was used. cDNAs were generated using the ProtoScript II First Strand cDNA Synthesis Kit and Oligo(dT) primers (*New England Biolabs*). RT-qPCR were performed on the Light Cycler 480II (*Roche*) with the primers listed in Supplementary Data 6 using *Eef1a1* and *Hprt* as housekeeping controls.

### RNA-seq and scRNA-seq

Total RNA from WT and dCD EpiLC (2 replicates) and EpiSC (2 replicates) were purified as described above. Bulk RNA-seq were generated following the protocol of the TruSeq stranded kit (*Illumina*).

For scRNA-seq, EpiSC were dissociated with Accutase and all other cell types with TripleExpress. Cells were then centrifuged and resuspended in PBS containing 0.04% BSA. Next, cells were passed through a strainer, the cell concentration was determined and the scRNA-seq libraries were prepared using the Chromium Single Cell Gene Expression (*10x Genomics*) according to the Single Cell 3’ Reagents Kit (v2) protocol for the single-cell experiment of the PGCLC differentiation (Fig. 1) and the Single Cell 3’ Reagents Kit (v3) protocol for the d4 EB from R1 WT and dCD cells (Fig. 6).

### Western Blot

Nuclei were isolated by incubating cells with lysis buffer (20 mM Tris pH 7.6, 100 mM NaCl, 300 mM sucrose, 3 mM MgCl_2_) containing freshly added protease inhibitors for 10 min at 4°C and then centrifuged for 10 min at 4°C and 3000 rpm. The resulting pellets, containing the cell nuclei, were treated with a high salt buffer (20 mM Tris pH 8.0, 400 mM NaCl, 2 mM EDTA pH 8.0) and disrupted with a glass homogenizer on ice. After incubation on ice for 30 min and centrifugation (24000 × g for 20 min at 4°C), supernatants were collected and protein concentration was estimated by a BCA-Assay. 20 µg of the resulting protein extracts were heated in Laemmli buffer at 95 °C for 5 min, loaded on 4–15% Mini-PROTEAN® TGX™ Precast Protein Gels (*Bio-Rad*) and transferred (190 mM glycine, 25 mM Tris, 20% Methanol, 0.1% SDS) to a PVDF membrane. After blocking with 5 % milk, the primary antibody (Supplementary Data 6) was incubated overnight at 4°C and the secondary antibody (coupled to a horseradish peroxidase (HRP)) for 1h at RT with washes in between. Finally, proteins were visualized using the lumi-light plus western blotting substrate (*Roche*).

### scRNA-seq data processing

The 10x Genomics scRNA-seq data generated in this study across different stages of PGCLC differentiation can be explored by opening the *.cloupe* file available through GEO (https://www.ncbi.nlm.nih.gov/geo/query/acc.cgi?acc=GSE155088) with the Loupe Cell Browser (https://www.10xgenomics.com). UMIs were counted using NCBI:GCA_000001635.6 and *cellranger-2.1.0*^87^. The resulting UMI values were aggregated into a single matrix with default normalization (‘‘–normalize=mapped’’) (Supplementary Data 1).

For the scRNA-seq data from E4.5 - E6.5 mouse embryos^43^ the count matrix (GSE121650) was normalized to FPKM (*edgeR*, Ensembl gene annotation, v87). Then, the previously generated lineage assignments^43^ were used to solely select epiblast cells for further analysis.

### scRNA-seq data analysis

The code used to define PGCLC genes is available through Github (https://github.com/ToreBle/Germline_competence). Briefly, *monocle2*^88^ was used to evaluate the *in vitro* scRNA-seq data generated across the different PGCLC differentiation stages. Therefore, k-means clustering was performed on the t-SNE plots (with k=3 for d2 EB and k=4 for d4 EB). From the resulting clusters, those containing PGCLC were identified by the enrichment of previously defined core PGC genes from d4/d6 PGCLC and E9.5 PGCs^89^. To determine the cellular identity of the remaining clusters found within the EB, the expression of lineage specific markers identified in E8.25 mouse embryos^42^ was used. To this end, all markers with a log2FoldChange >2.5 were considered. Each EB cluster was annotated as equivalent to the mouse embryonic tissue for which we observed the most significant enrichment in the expression of the corresponding marker genes.

The ESC, EpiLC, EpiSC and PGCLC gene sets were defined by differential expression using *Seurat*^90^ and the “negbinom” option for differential expression testing. ESC, EpiLC and EpiSC genes were determined by differential expression between the stage indicated vs the remaining two (e.g. ESC vs. d2 EpiLC and EpiSC) From this analysis only the genes upregulated in the particular stage (adjusted p-value < 0.005), a high expression in the stage of interest (more than 20 %, pct.1 >0.2) and a low expression distribution in the others (less than 40%, pct.2<0.4) were considered. Similar PGCLC genes were determined by differential expression between the d2+d4 PGCLC clusters and the remaining clusters from d2 and d4 EB. Again, from this analysis only the genes upregulated in PGCLC (adjusted p-value < 0.005) and with high expression in PGCLC (expressed in 20 % of the PGCLC) and a low expression distribution in the other analyzed cells (expressed in less than 40% of non-PGCLC EB cells) were considered. In the case of PGCLC genes this resulted in 389 PGCLC genes (Supplementary Data 2). To quantify the expression of PGCLC genes in different analyses, the UMI count matrix (of the *in vitro* stages) or FPKM-normalized data (of the *in vivo* stages) were used to calculate the mean expression of all PGCLC genes within a cell or the mean expression of each PGCLC gene across all cells of a stage.

For the RNA velocity analysis (Supplementary Fig. 1e), spliced and unspliced read counts were obtained with *kallisto*^91^ and *bustools*^92^, parsed into R-3.6.1 to create a *Seurat* object and t-SNE plot which was then overlaid by the RNA velocity calculations from *velocyto.R*^93^.

The estimation of transcriptional noise for the epiblast stages *in vivo* and *in vitro* was performed like in^44^. First, the 500 most variable genes for each stage were selected and pairwise compared by Spearman correlations. Then the Spearman correlation values were transformed into distance measurements (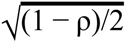) that were considered to represent transcriptional noise.

For the single-cell RNA-seq data set of the d4 EB from R1 WT and dCD cells, single cells with a high percentage of mitochondrial gene expression were discarded, by considering only single cells with at least 2000 expressed genes. The clusters (Supplementary Fig. 6e) were identified in the combined dataset of R1 WT and dCD cells by shared nearest neighbor (SNN) modularity optimization based clustering algorithm from Seurat with a resolution of 0.3. This resulted in 9 clusters that were classified based on the marker gene expression of E8.25 mouse embryos^42^. Those clusters that did not show any specific marker gene expression from this cluster (e.g. 2-cell-like) were identified by differentially expressed genes within the cluster. The *Prdm1orDppa3+/Klf4-* cells were defined using the following gene expression values; Dppa3 > 0.1 OR Prdm1>1 AND Klf4<1. The single-cell expression correlation values were determined by the spearman correlation of the top 1000 most variable genes in each cluster and the correlation of their expression among the cells belonging to the same cluster.

### ChIP-seq data processing

For ChIP-seq data processing, single end reads were mapped to the mouse genome (*mm10*) using *BWA*^94^. After duplication removal with the *MarkDuplicates* function from the *Picard* tools (http://broadinstitute.github.io/picard/), reads within blacklisted regions (https://www.encodeproject.org/annotations/ENCSR636HFF/) were discarded and the aligned reads were normalized with *deeptools-3.3.1*^95^ to 1x sequencing depth (as RPGC: Reads per genomic content). An overview of all the ChIP-seq experiments performed in this study is listed in Supplementary Data 6 and, briefly, it includes the following datasets:

- H3K4me1 ChIP-seq experiments in ESC (n = 2), EpiLC (n = 4) and EpiSC (n = 2) performed in R1 and E14Tg2a cell lines.
- H3K4me2 ChIP-seq experiments in ESC (n = 2), EpiLC (n = 3) and EpiSC (n = 3) performed in R1 and E14Tg2a cell lines.
- H3K4me3 ChIP-seq experiments in ESC (n = 2), EpiLC (n = 2) and EpiSC (n = 2) performed in R1 and E14Tg2a cell lines.
- H3K27ac ChIP-seq experiments in ESC (n = 2), EpiLC (n = 4) and EpiSC (n=2) performed in R1 and E14Tg2a cell lines.
- NANOG-HA ChIP-seq experiments in EpiSC were performed as two biological replicates in E14Tg2a.
- Additional ChIP-seq experiments were performed as single replicates (Supplementary Data 6).

All the processed ChIP-seq data can be explored using the following UCSC browser sessions:

- Comparison of the pluripotent stages (related to Fig. 2-3): http://genome-euro.ucsc.edu/s/Tore/Comparison%20of%20the%20pluripotent%20stages
- PRDM14/NANOG overexpression (related to Fig. 4): http://genome-euro.ucsc.edu/cgi-bin/hgTracks?hgS_doOtherUser=submit&hgS_otherUserName=Tore&hgS_otherUserSessionName=PRDM14%2FNANOG%20overexpression
- H3K4me1/2 deficient cells and *Otx2*^−/−^ cells (related to Fig. 4-5): http://genome-euro.ucsc.edu/cgi-bin/hgTracks?hgS_doOtherUser=submit&hgS_otherUserName=Tore&hgS_otherUserSessionName=H3K4me1%2F2%20deficiency%20and%20Otx2%2D%2F%2D

### ATAC-seq data processing

ATAC-seq experiments were performed as two biological replicates for each of the investigated cell types (i.e. ESC, EpiLC, EpiSC). For ATAC-seq data processing, paired-end reads were mapped to the mouse genome (*mm10*) using *BWA*^94^. Read duplicates and reads within blacklisted regions were discarded. Given the concordance of the ATAC-seq replicas (Spearman correlation of a 2 kb window for ESC: 0.86; EpiLC: 0.88; EpiSC: 0.86), BAM files for each stage were merged and converted into bigWig files by normalization to 1x sequencing depth with *deeptools-3.3.1*^95^. The processed ATAC-seq data can be also found in the UCSC browser sessions mentioned above.

### Enhancer definition

The code used to define enhancers is available through Github (https://github.com/ToreBle/Germline_competence).

PGCLC enhancers: The H3K27ac ChIP-seq data (SOLiD sequencing) from d2 and d6 sorted PGCLC^15^ were aligned with the default settings of *novoalignCS* (V1.06.09, *Novocraft Technologies*). For visualization, replicate BAM files for each stage were merged and normalized to 1 × sequencing depth. Briefly, H3K27ac peaks were called from both replicates of d6 PGCLC with MACS2^96^ using broad settings (--broad -m 5 50 --fix-bimodal --extsize 200) and q-values < 1 × 10^−3^. Next, all peaks within a distance of 1kb from both replicates d6 PGCLC were merged with *bedtools* (https://github.com/arq5x/bedtools2). The resulting regions were subtracted from blacklisted and promoter regions (−/+ 2 kb of the Ensembl gene annotation, v86) with *bedtools* (Supplementary Fig. 2a). PGCLC enhancers were defined as the subset of the distal H3K27ac peaks identified in d6 PGCLC that could be physically linked to PGCLC genes according to capture HiC data generated in ESC by^47^ (Supplementary Data 1). Thereby, distal d6 PGCLC H3K27ac peaks and PGCLC genes were linked if the two anchors of a Capture HiC interaction occurred within 1 Kb of a d6 PGCLC H3K27ac peak and 3 Kb of the PGCLC gene TSS.

For the statistical comparisons of epigenetic signals within PGCLC enhancers in different cell types, we first determined the average signals of the ChIP-seq, ATAC-seq or genome-wide CpG bisulfite sequencing within −/+ 1kb of the PGCLC enhancers in d2 EpiLC and EpiSC using *deeptools-3.3.1*^95^. Then, the effect size of paired wilcoxon tests were calculated by dividing the z-statistics by the square roots of the sample sizes using *rstatix* and *boots.ci* for the approximation of the confidence intervals (confidence level: 0.95).

EpiLC/EpiSC enhancers: Briefly, H3K27ac peaks were called in both cell types separately with MACS2^96^ using broad settings (--broad -m 5 50 --fix-bimodal --extsize 200) and q-values < 1 × 10^−3^. The resulting regions were subtracted from blacklisted and promoter regions (−/+ 2 kb of the Ensembl gene annotation, v86) using *bedtools*. EpiLC and EpiSC enhancers were obtained from these regions by selecting only H3K27ac peaks located within 500 kb of the gene sets defined by single-cell RNA-seq for each cell type. The enhancer assignment to the proximal genes was done with *GREAT-4.0.4*^97^ and only enhancers with a minimum distance of 3.5 kb to the nearest transcription start site (TSS) were considered.

### CpG methylation analysis

The analysis of the local bisulfite sequencing experiments performed for selected PGCLC enhancers was evaluated with BISMA^98^.

Genome-wide DNA methylation data was analysed with *Bismark*^99^. However, as the considered data sets were prepared with slightly different protocols, the preprocessing steps were adjusted accordingly: For the whole-genome bisulfite sequencing data from 2i ESC (GSE41923), d2 EpiLC and EpiSC (GSE70355), the adapter trimming was performed with *Trim Galore* (http://www.bioinformatics.babraham.ac.uk/projects/trim_galore/) using the default settings; for data sets generated by post bisulfite adaptor tagging (pbat), either 9 (data from d2 EpiLC and PGCLC (DRA003471)) or 6 bp (genome-wide methylation data generated in this study) were removed. The DNA methylation data from E4.5 - E6.5 epiblasts were generated by STEM-seq (GSE76505), and, in this case, adapter sequences were removed with *cutadapt*^100^ and *Trim Galore,* respectively. For all the previous samples, reads were mapped with *Bismark-v0.16.1*^99^ and *bowtie2-2.2.9*^101^, using the “-pbat” setting for the STEM-seq and pbat samples. For paired-end pbat samples the unmapped reads were remapped as single-end reads. Then, for each cell type all available datasets were combined to estimate the CpG methylation levels with the *Bismark methylation extractor*. Finally only CpGs with a coverage of 3 - 100 reads were considered.

### CpG methylation heterogeneity and scNMT-seq analysis

With the scNMT-seq method the transcriptome, methylome (CpG methylation) and chromatin accessibility (GpC methylation) are recorded from the same single cell^102^. Among all the single cells analyzed in E4.5, E5.5 and E6.5 mouse embryos by^43^, we only considered those assigned to the epiblast by scRNA-seq (https://github.com/rargelaguet/scnmt_gastrulation). Then, epiblast cells were additionally filtered according to their methylome (GSE121690) and only cells with a CpG coverage > 10^2^ at PGCLC enhancers were considered. This resulted in 258 epiblast cells from E4.5, E5.5 and E6.5 with high quality single cell CpG methylation data.

The single cell CpG methylation for each cell was stored in a bedGraph file. From each bedGraph file the genome-wide CpG methylation levels of individual cells were determined as the mean methylation of all covered CpGs in each single cell. For the analysis of mCpG levels and CpG coverage within PGCLC enhancer Group I, the genome-wide bedGraph files were subtracted to obtain bedGraph files with the PGCLC enhancer regions (Supplementary Data 3) using *bedtools* (https://github.com/arq5x/bedtools2) from which the mean methylation of all covered CpGs was determined in each single cell.

CpG methylation heterogeneity was estimated with the *PDclust* package^66^. The number of CpGs covered in each pair of cells resulted in approximately 150 CpGs for each pairwise comparison when PGCLC enhancers were considered. Then, the average of the absolute difference in the methylation values for all the CpGs covered for each pairwise comparison were computed as a dissimilarity matrix.

For the scRNA-seq data, the mean expression of all PGCLC genes (FPKM-normalized) linked to Group I PGCLC enhancers was determined per cell.

### Bulk RNA-sequencing

Public available RNA-seq data generated across different time-points of EpiLC differentiation ^103^, *Otx2*^−/−^ EpiLC^53^ and *Prdm14*^−/−^ EpiLC^14^ were mapped with *STAR*^104^ to the mouse reference genome (Ensembl gene annotation, v99) and reads within genes were counted with *featureCounts*^105^. The rlog normalization was generated with *DESeq2*^106^.

For the bulk RNA-seq data generated in R1 WT and dCD cells, we analyzed three replicates for R1 WT and the dCD ESC previously generated by Dorighi et al., 2017, two replicates for R1 WT and dCD d2 EpiLC and two replicates for R1 WT and dCD EpiSC. The transcript abundance was determined with the bootstrap mode of *kallisto* (-b 100)^91^ and the differential expression analyses were performed with *sleuth* on the gene-level (Ensembl gene annotation, v96)^91, 107^.

## Supplementary Data

Supplementary Data 1 - UMI count matrix of all cells analyzed across different stages of PGCLC differentiation

Supplementary Data 2 - ESC, EpiLC, EpiSC and PGCLC gene sets based on single-cell RNA-seq profiling across different stages of PGCLC differentiation

Supplementary Data 3 - EpiLC, EpiSC and PGCLC enhancers coordinates and linked genes

Supplementary Data 4 - RNA-seq results in WT vs Mll3/4 dCD cells

Supplementary Data 5 - UMI count matrix of all cells analyzed from d4 EB (WT & dCD)

Supplementary Data 6 - Resources

## Supplemental Figures

**Supplementary Fig.1:**
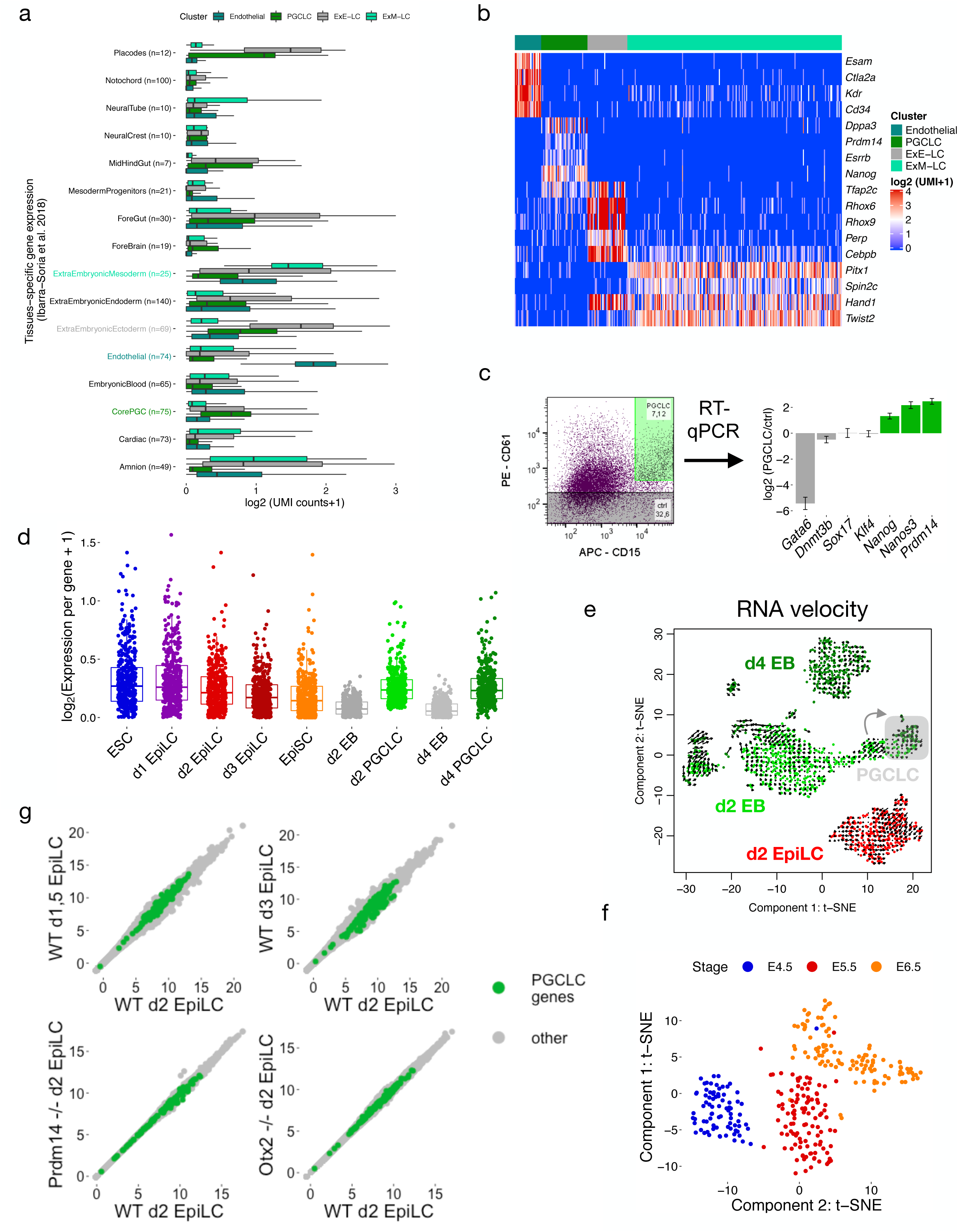
Characterization of the PGCLC differentiation system by scRNA-seq (related to Fig. 1). a.) The average gene expression for the specific markers defining the main embryonic and extraembryonic tissues found within E8.25 mouse embryos (Ibarra-Soria *et al.* 2018) is shown for the cell clusters identified within d4 EBs. Since PGC markers were not identified by Ibarra-Soria *et al.* 2018, we used the core PGC gene set previously defined by Nakaki *et al*. 2016 as a representative set of PGC markers expressed in E9.5 embryos. The numbers next to each tissue name indicates the genes considered as specific markers for each tissue (tissue markers with logFC >2.5). b.) Heatmap showing the expression of representative markers for the main four cell clusters identified within d4 EBs (Fig. 1C; Endothelial, PGCLC, ExE-LC and ExM-LC). The expression values are displayed as UMI (unique molecular identifier) counts in log2 scale. c.) D4 EB cells were sorted by FACS into CD15^+^CD61^+^ (*i.e.* PGCLC) or CD61^−^ (*i.e.* non-PGCLC control cells). Then, the expression of the indicated genes, including the PGC markers *Prdm14*, *Nanos3* and *Nanog,* was determined by RT-qPCR in those two cell populations. The error bars represent standard deviations from three technical replicates. d.) Expression dynamics of the PGCLC genes during PGCLC formation. Each dot represents the mean expression of a PGCLC gene in all cells belonging to the indicated stages. EB refers to any cell of the EB except those identified as PGCLC. e.) RNA velocity was used as a computational approach to investigate potential transcriptional priming of the PGCLC expression program in d2 EpiLC. The RNA velocity analysis, which takes into account unspliced transcript reads for lineage tracing, did not show any evidence of transcriptional priming of d2 EpiLC (red) towards the PGCLC cluster (gray). In contrast, this same analysis revealed that a fraction of the cells profiled for the d2 EB (with a mixed mesodermal/PGCLC identity; gray arrow) seem to acquire a transcriptional program similar to the one found for PGCLC (shadowed in gray), indicating the feasibility of the RNA velocity method to detect transcriptional priming. f.) t-SNE plot of the *in vivo* single-cell RNA-seq data from E4.5, E5.5 and E6.5 (Argelaguet *et al.* 2019). Only cells determined as epiblast cells by Argelaguet *et al.* 2019 are considered in the plot. g.) Scatter plots comparing the transcriptomes of WT d2 EpiLC (x-axis) with those of d1.5 EpiLC, d3 EpiLC, *Prdm14*^−/−^ d2 EpiLC or *Otx2*^−/−^ d2 EpiLC (y-axis). All genes considered as expressed are shown as dots, with the PGCLC genes highlighted in green. The gene expression values are r-log normalized. The RNA-seq data was obtained from Yang *et al*. 2019, Buecker *et al*. 2014 and Shirane *et al*. 2016.

**Supplementary Fig. 2:**
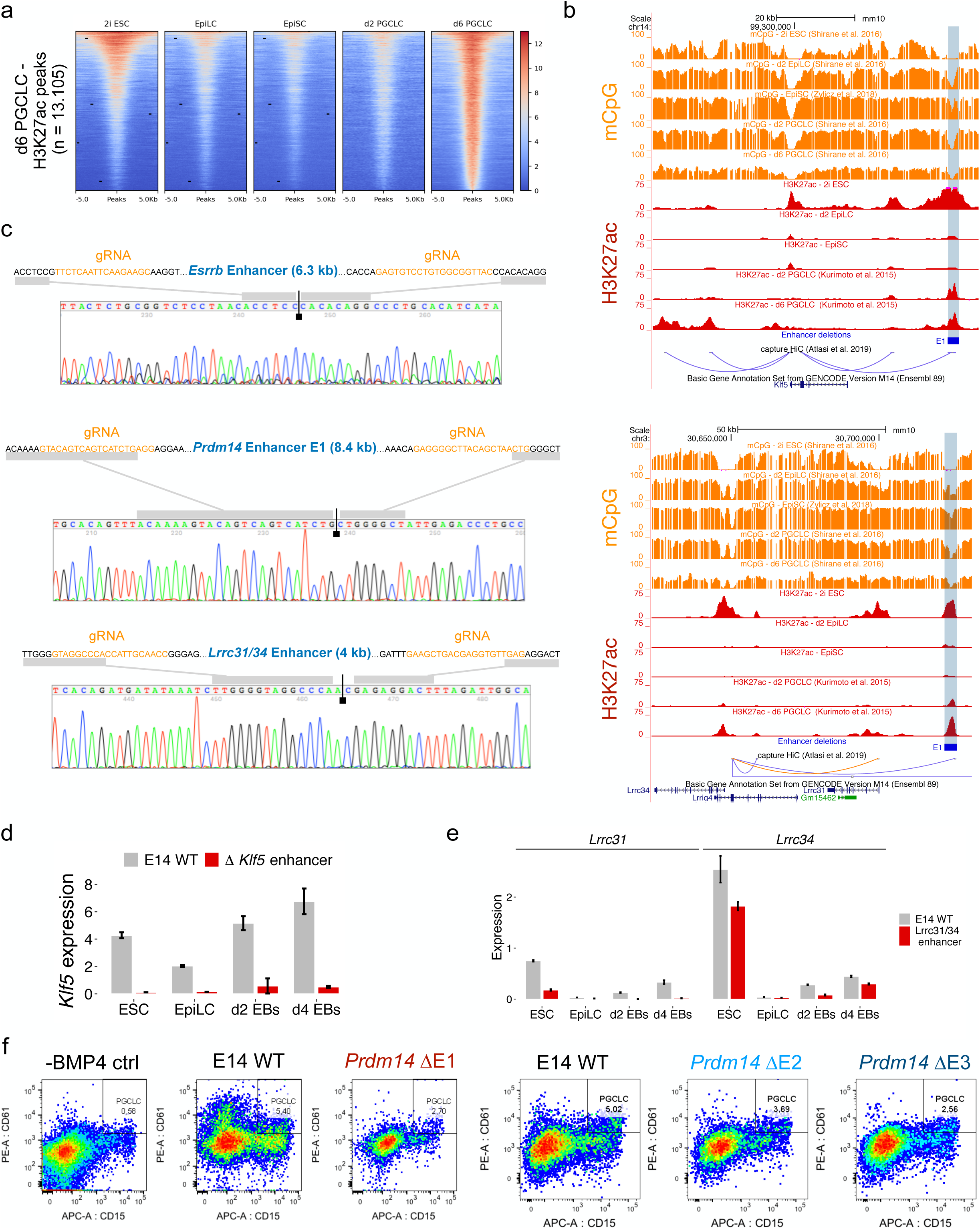
Deletion and functional evaluation of representative PGCLC enhancers (related to Fig. 2). a.) Heatmaps showing H3K27ac ChIP-seq signals during PGCLC differentiation around all the distal H3K27ac peaks called in d6 PGCLC. The H3K27ac peaks were called using two H3K27ac ChIP-seq replicates generated in d6 PGCLC and only those peaks located >2 Kb from gene TSS were considered (see Methods). The H3K27ac ChIP-seq data from d2 and d6 PGCLC was obtained from Kurimoto *et al*. 2015. b.) Genome-browser view of the H3K27ac and CpG methylation dynamics during PGCLC differentiation for the PGCLC enhancers, highlighted in blue, found within within the *Klf5* (top) and *Lrrc31/Lrrc34* (bottom) loci. The blue rectangles denote the enhancer deletions generated in mESC using CRISPR/Cas9 technology. According to capture Hi-C data generated in ESC (Atlasi *et al.* 2019), the *Klf5* enhancer is physically linked to the *Klf5* TSS, while the *Lrrc31/ Lrrc34* enhancer is directly linked to the *Lrrc34* TSS, which itself interacts with *Lrrc31* (orange line). The H3K27ac PGCLC data was obtained from Kurimoto *et al.* 2015 and the CpG methylation data from Shirane *et al.* 2016. c.) The different PGCLC enhancer deletions generated in ESC were confirmed by PCR genotyping followed by Sanger sequencing. This is illustrated by chromatograms showing the deletions generated for the *Esrrb*, *Prdm14* E1 and *Lrrc31/34* PGCLC enhancers. The reference sequence for each enhancer together with the corresponding CRISPR/Cas9 gRNAs (highlighted in orange) are shown above the chromatograms. d-e.) *Klf5* (d)*, Lrrc31* and *Lrrc34* (e) expression were measured by RT-qPCR in d4 EB differentiated from WT ESC and two different ESC clonal lines with the *Lrrc31/Lrrc34* (d) or *Klf5* (e) enhancer deletions. The expression values were normalized to two housekeeping genes (*Eef1a1* and *Hprt*). Error bars represent standard deviations from 6 measurements (two clonal lines × three technical replicates). f.) Representative examples of the PGCLC quantifications performed by FACS after four days of PGCLC differentiation using WT ESC (E14 WT) or ESC with deletions of each of the three *Prdm14* enhancers (ΔE1-E3). As a negative control, the WT ESC were also differentiated in the absence of BMP4 (-BMP4 ctrl). PGCLC were defined as the CD15^+^CD61^+^ cells found within d4 EB.

**Supplementary Fig. 3:**
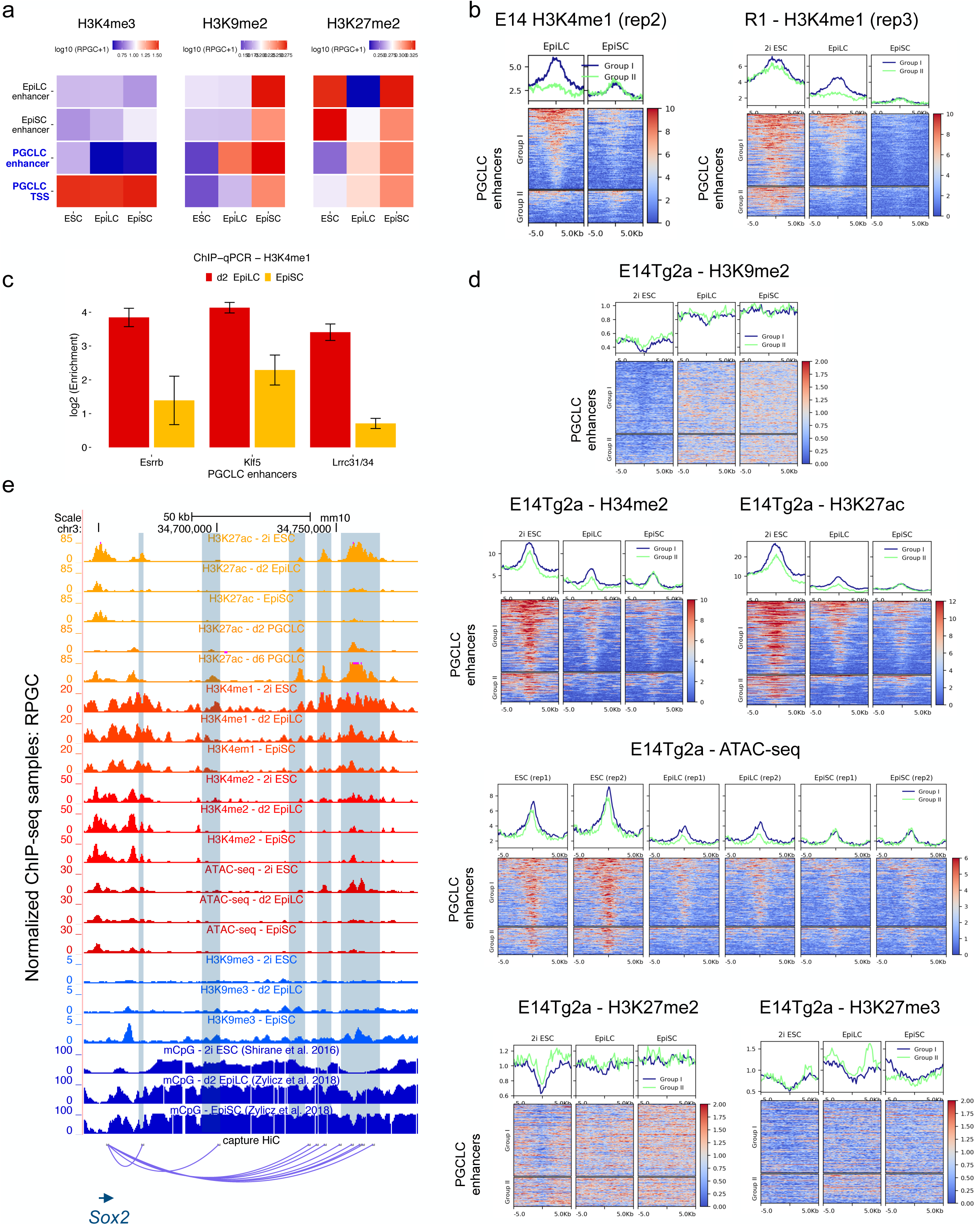
Epigenomic profiling of PGCLC enhancers (related to Fig. 3). a.) Summary plot showing the average levels of H3K4me3, H3K9me2 and H3K27me2 measured in 2i ESC, d2 EpiLC and EpiSC for the EpiLC, EpiSC and PGCLC enhancers as well as the TSS of the PGCLC genes. Quantifications were performed by measuring the average signals of each epigenetic mark within −/+ 1kb of the enhancers orTSS. RPGC: Reads per genomic content. b.) Average profile (top) and heatmap (bottom) plots showing H3K4me1 levels for the Group I and Group II PGCLC enhancers in 2i ESC, d2 EpiLC and EpiSC. The H3K4me1 signals correspond to the second (E14 cells) and third (R1 cells) ChIP-seq replicates generated in each cell type. c.) H3K4me1 ChIP-qPCR analyses for the PGCLC enhancers of *Esrrb*, *Klf5* and *Lrrc31/34* in d2 EpiLC and EpiSC differentiated from E14 ESC. The H3K4me1 enrichments levels were normalized to a negative control region and are shown in log2 scale. d.) Average profile (top) and heatmap (bottom) plots showing H3K9me2, H3K4me2, H3K27ac and ATAC-seq (from two biological replicates), H3K27me2 and H3K27me3 levels for the Group I and Group II PGCLC enhancers in 2i ESC, d2 EpiLC and EpiSC. e.) Genome-browser view showing representative PGCLC enhancers interacting with *Sox2* according to capture Hi-C generated in ESC (Atlasi *et al.* 2019, blue arc) as well as the levels of several epigenetic marks in 2i ESC, d2 EpiLC and EpiSC. The H3K27ac PGCLC data was obtained from Kurimoto *et al.* 2015 and the CpG methylation data from Shirane *et al.* 2016 and Zylicz *et al.* 2015.

**Supplementary Fig. 4:**
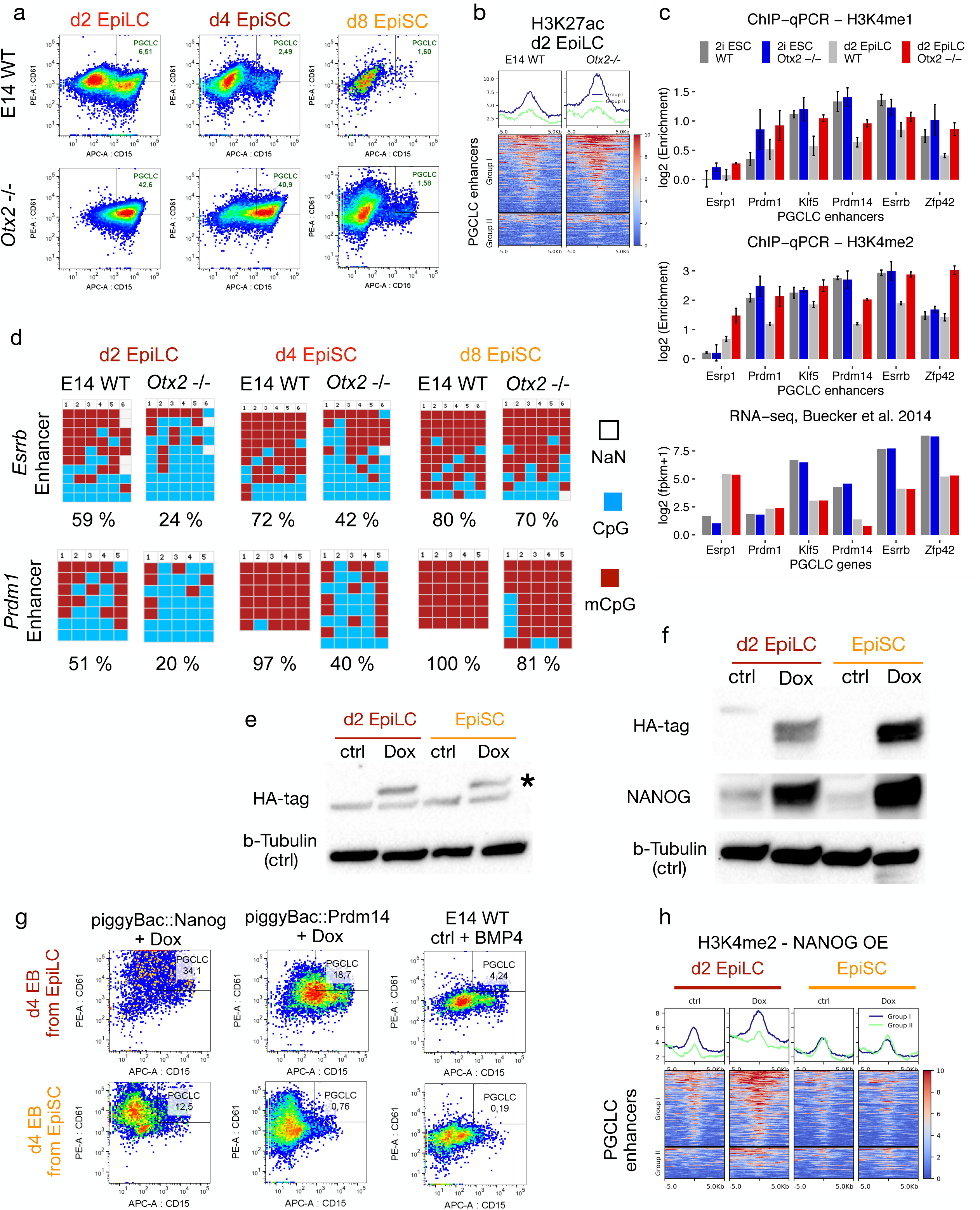
Characterization of PGCLC enhancers in *Otx2*^−/−^ cells and upon overexpression of major germline regulators (related to Fig. 4). a.) Representative examples of the PGCLC quantifications performed by FACS after four days of PGCLC differentiation starting from d2 EpiLC, d4 EpiSC and d8 EpiSC for either E14 WT (top) or E14 *Otx2*^−/−^ (bottom) cells. PGCLC were quantified as the CD15^+^CD61^+^ cells found within d4 EB. b.) Average profile (top) and heatmap (bottom) plots showing H3K27ac signals for the Group I and Group II PGCLC enhancers in E14 WT and *Otx2*^−/−^ d2 EpiLC. c.) H3K4me1 (top) and H3K4me2 (middle) ChIP-qPCR analyses for selected PGCLC enhancers in E14 WT and *Otx2*^−/−^ cells (2i ESC and d2 EpiLC). H3K4me1 and H3K4me2 enrichments levels were normalized to a negative control region and shown in log2 scale. The bottom panel shows the expression values for the genes associated with the selected PGCLC enhancers using RNA-seq data from Buecker et al. 2014. d.) CpG methylation levels of selected PGCLC enhancers linked to *Esrrb* and *Prdm1* genes were measured by bisulfite sequencing in d2 EpiLC, d4 EpiSC and d8 EpiSC differentiated from E14 WT and *Otx2*^−/−^ ESC. The columns of the plots correspond to individual CpG dinucleotides located within the indicated enhancer. Unmethylated CpGs are shown in blue, methylated CpGs in red and CpGs that were not sequenced are shown in gray. The rows represent the sequenced alleles in each cell line. e-f.) Western blot analysis of the inducible overexpression of exogenous PRDM14-HA (e) and NANOG-HA (f). EpiLC or EpiSC were either left untreated (ctrl) or treated with Dox for 18 hours. (e) PRDM14-HA levels were measured using an anti-HA antibody and the band corresponding to PRDM14-HA is indicated with an asterisk. (f) NANOG levels were measured using anti-HA or anti-NANOG antibodies. B-Tubulin was used as a loading control. g.) ESC lines enabling the inducible overexpression of exogenous NANOG-HA (piggyBac::Nanog) or PRMD14-HA (piggyBac::Prdm14) were differentiated into d2 EpiLC and EpiSC. Next, the resulting d2 EpiLC or EpiSC were differentiated into PGCLC upon overexpression (+Dox) of NANOG-HA or PRDM14-HA and in the absence of growth factors. Similarly, E14 WT d2 EpiLC and EpiSC were also differentiated into PGCLC in the presence of growth factors (ctrl + BMP4). Representative examples of the PGCLC quantifications performed by FACS after four days of PGCLC differentiation are shown. PGCLC were quantified as the percentage of CD15+ CD61+ cells found within d4 EB. h.) The ESC line enabling the inducible overexpression of exogenous NANOG-HA was differentiated into d2 EpiLC and EpiSC. Next, the cells either left untreated (ctrl) or treated with Dox for 18 hours and H3K4me2 ChIP-seq experiments were performed. The average profile (top) and heatmap (bottom) plots show H3K4me2 levels within Group I and Group II PGCLC enhancers.

**Supplementary Fig. 5:**
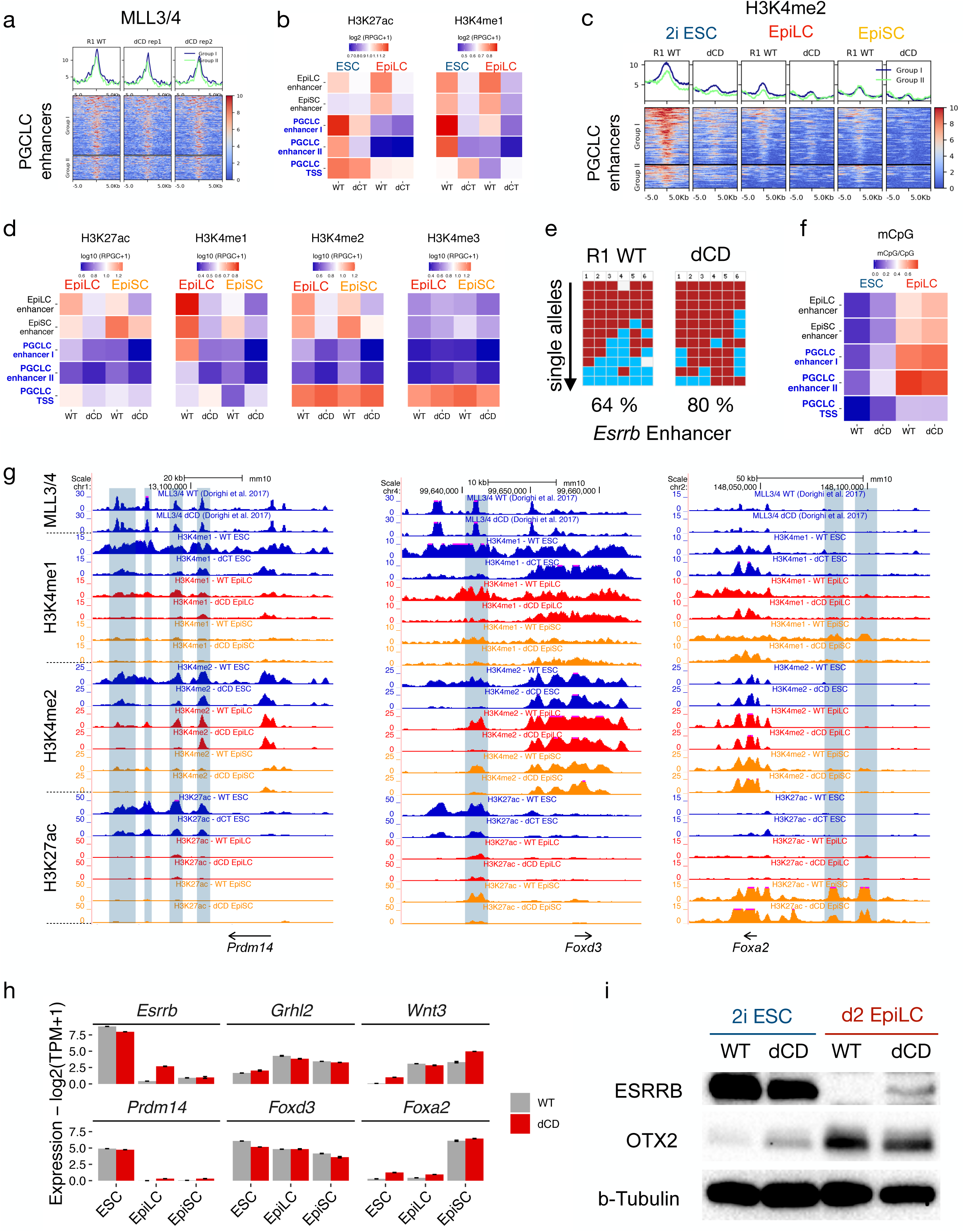
Epigenomic and transcriptional characterization of MLL3/4 catalytic mutant cells (related to Fig. 5). a.) Average profile (top) and heatmap (bottom) plots showing the binding of WT and dCD MLL3/4 proteins for the Group I and II PGCLC enhancers in 2i ESC (ChIP-seq data from Dorighi *et al*. 2017). The WT MLL3/4 ChIP-seq experiments were merged from two replicates. b.) Summary plot showing the average H3K27ac and H3K4me1 levels in WT and dCT cells (ESC and d2 EpiLC) for the EpiLC, EpiSC and PGCLC enhancers as well as the TSS of the PGCLC genes. Quantifications were performed by measuring the average signals of each epigenetic mark within −/+ 1kb of the enhancers/TSS. RPGC: Reads per genomic content. c.) Average profile (top) and heatmap (bottom) plots showing H3K4me2 levels for the Group I and Group II PGCLC enhancers in R1 WT and dCD ESC lines as well as upon their differentiation into d2 EpiLC and EpiSC. d.) Summary plot showing the average H3K27ac, H3K4me1, H3K4me2 and H3K4me3 levels in WT and dCD cells (d2 EpiLC and EpiSC) for the EpiLC, EpiSC and PGCLC enhancers as well as the TSS of the PGCLC genes. Quantifications were performed by measuring the average signals of each epigenetic mark within −/+ 1kb of the enhancers orTSS. The H3K4me1 ChIP-seq data shown for WT d2 EpiLC and EpiSC are the same ones used in Fig. 3c as third replicates. RPGC: Reads per genomic content. e.) The CpG methylation levels of the *Esrrb* enhancer were determined by bisulfite sequencing in R1 WT and dCD d2 EpiLC. The columns of the plots correspond to individual CpG dinucleotides located within the *Esrrb* enhancer. Unmethylated CpGs are shown in blue, methylated CpGs in red and CpGs that were not sequenced are shown in gray. The rows represent the sequenced alleles in each cell line. f.) Summary plot showing the average ratio of methylated CpG in WT and dCD cells (2i ESC and d2 EpiLC) for the EpiLC, EpiSC and PGCLC enhancers as well as the TSS of the PGCLC genes. Quantifications were performed by measuring the average signals of each epigenetic mark within −/+ 1kb of the enhancers orTSS. g.) Genome browser view showing H3K4me1, H3K4me2 and H3K27ac profiles in both WT and MLL3/4 catalytic mutant cells (ESC, EpiLC and EpiSC) around representative PGCLC (*Prdm14)*, EpiLC (*Foxd3*) and EpiSC (*Foxa2*) genes and their associated enhancers. h.) Expression levels, as measured by RNA-seq, for the representative PGCLC, EpiLC and EpiSC genes shown in Fig. 5c and Supplementary Fig. 5g in WT (gray) and dCD (red) ESC cells as well as upon their differentiation into EpiLC and EpiSC. The RNA-seq data in ESC was obtained from Dorighi et al. 2017 and the experiments in d2 EpiLC and EpiSC were performed in duplicates. i.) Western Blot analysis of the ESRRB and OTX2 protein levels present in WT and dCD ESC as well as upon their differentiation into d2 EpiLC. B-Tubulin was used as a loading control.

**Supplementary Fig. 6:**
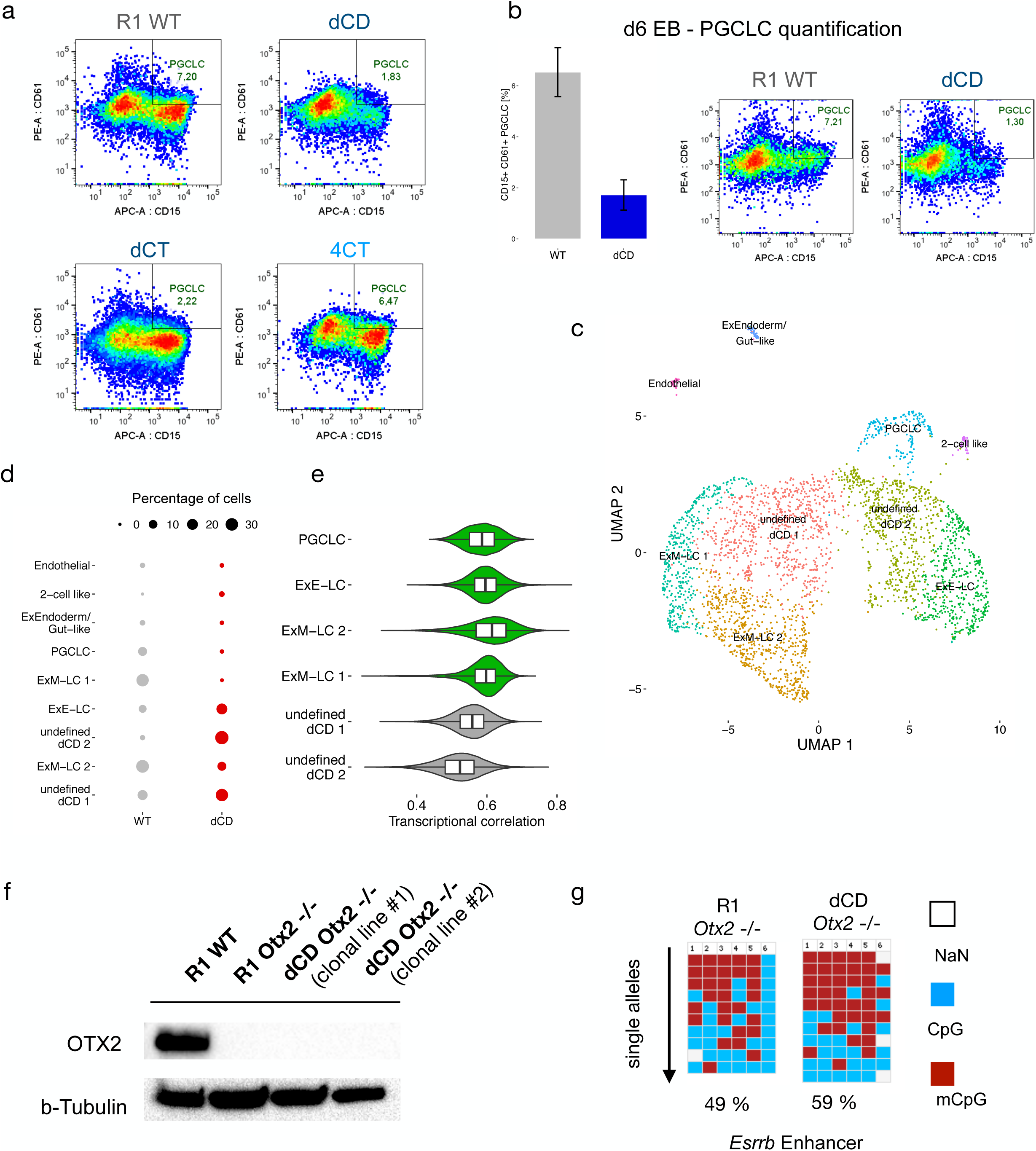
Evaluation of the importance of H3K4me1 during PGCLC differentiation (related to Fig. 6). a.) Representative examples of the PGCLC quantifications performed by FACS after four days of PGCLC differentiation using WT ESC (R1 WT) or with the indicated MLL3/4 catalytic mutant ESC lines. PGCLC were quantified as the CD15^+^ CD61^+^ cells found within d4 EB. b.) WT ESC and *Mll3/4* dCD ESC lines were differentiated into PGCLC and PGCLC were quantified in d6 instead of d4 EB. Examples are shown on the left and the PGCLC quantification of CD15^+^ CD61^+^ cells within d6 EB from two biological replicates are shown on the right. c.) UMAP plot showing the different cell clusters identified within WT and dCD d4 EB according to the single-cell RNA-seq data. The single-cell RNA-seq data was subject to clustering analysis (Shared Nearest Neighbor) and the cellular identity of the resulting clusters was determined by differential expression analysis between clusters as well as by using specific markers of the main embryonic and extraembryonic tissues found within E8.25 mouse embryos (Ibarra-Soria *et al.* 2018) (Fig. S1a). d.) Percentage of WT and dCD cells present within each of the clusters identified in d4 EB as described in (c). e.) Transcriptional correlation of the main d4 EB clusters as described in (c-d). Spearman correlations of the 1000 most variable genes between single cells of the indicated clusters are shown. f.) Western blot analysis of the OTX2 levels in *Otx2*^−/−^ and dCD *Otx2*^−/−^ d2 EpiLC. B-Tubulin was used as a loading control. g.) The CpG methylation levels of the *Esrrb* enhancer were determined by bisulfite sequencing in R1 *Otx2*^−/−^ and dCD *Otx2*^−/−^ d2 EpilC. The columns of the plots correspond to individual CpG dinucleotides located within the *Esrrb* enhancer. Unmethylated CpGs are shown in blue, methylated CpGs in red and CpGs that were not sequenced are shown in gray. The rows represent the sequenced alleles in each cell line.

**Supplementary Fig. 7:**
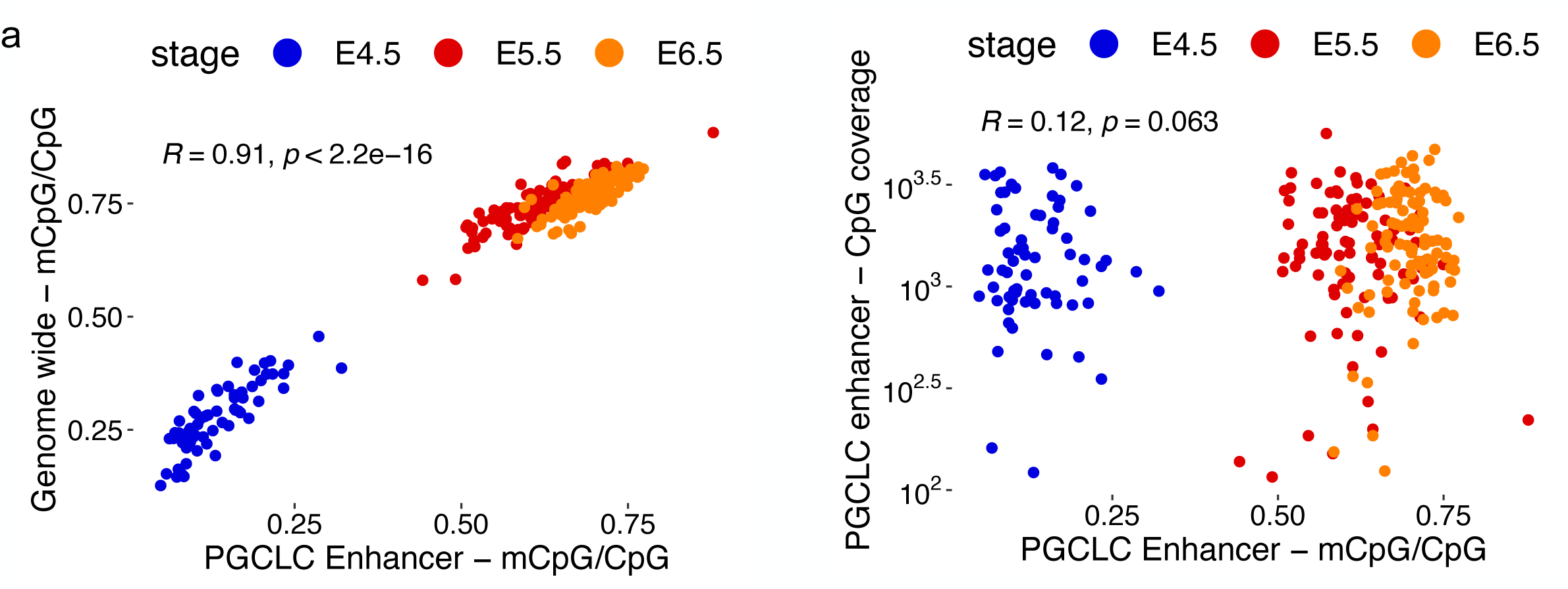
Analysis of PGCLC enhancers using *in vivo* single-cell CpG methylation data (related to Fig. 7). a.) Scatterplots showing the correlation between (left) genome-wide mCpG or (right) CpG coverage and the mean mCpG/CpG ratio of all PGCLC enhancers within individual cells belonging to E4.5, E5.5 and E6.5 epiblasts (n=261 cells). The correlation coefficient (R) and p-value were calculated using Spearman correlation.

## References

1. Waddington, C. H. Organisers and Genes by C. H. Waddington. (1940).

2. Solini, G. E., Dong, C. & Saha, M. Embryonic transplantation experiments: Past, present, and future. Trends Dev. Biol. 10, 13–30 (2017).

3. Ohinata, Y. et al. Blimp1 is a critical determinant of the germ cell lineage in mice. Nature 436, 207–213 (2005).

4. Ohinata, Y. et al. A signaling principle for the specification of the germ cell lineage in mice. Cell 137, 571–584 (2009).

5. Kalkan, T. et al. Tracking the embryonic stem cell transition from ground state pluripotency. Development 144, 1221–1234 (2017).

6. Mulas, C., Kalkan, T. & Smith, A. NODAL Secures Pluripotency upon Embryonic Stem Cell Progression from the Ground State. Stem Cell Reports 9, 77–91 (2017).

7. Smith, A. Formative pluripotency: the executive phase in a developmental continuum. Development vol. 144 365–373 (2017).

8. Bleckwehl, T. & Rada-Iglesias, A. Transcriptional and epigenetic control of germline competence and specification. Curr. Opin. Cell Biol. 61, 1–8 (2019).

9. Respuela, P. et al. Foxd3 Promotes Exit from Naive Pluripotency through Enhancer Decommissioning and Inhibits Germline Specification. Cell Stem Cell 18, 118–133 (2016).

10. Hayashi, K., Ohta, H., Kurimoto, K., Aramaki, S. & Saitou, M. Reconstitution of the mouse germ cell specification pathway in culture by pluripotent stem cells. Cell 146, 519–532 (2011).

11. Hackett, J. A. et al. Tracing the transitions from pluripotency to germ cell fate with CRISPR screening. Nat. Commun. 9, 4292 (2018).

12. Murakami, K. et al. NANOG alone induces germ cells in primed epiblast in vitro by activation of enhancers. Nature 529, 403–407 (2016).

13. Mitani, T. et al. Principles for the regulation of multiple developmental pathways by a versatile transcriptional factor, BLIMP1. Nucleic Acids Res. 45, 12152–12169 (2017).

14. Shirane, K. et al. Global Landscape and Regulatory Principles of DNA Methylation Reprogramming for Germ Cell Specification by Mouse Pluripotent Stem Cells. Dev. Cell 39, 87–103 (2016).

15. Kurimoto, K. et al. Quantitative Dynamics of Chromatin Remodeling during Germ Cell Specification from Mouse Embryonic Stem Cells. Cell Stem Cell 16, 517–532 (2015).

16. von Meyenn, F. et al. Comparative Principles of DNA Methylation Reprogramming during Human and Mouse In Vitro Primordial Germ Cell Specification. Dev. Cell 39, 104–115 (2016).

17. Saitou, M., Kagiwada, S. & Kurimoto, K. Epigenetic reprogramming in mouse pre-implantation development and primordial germ cells. Development vol. 139 15–31 (2012).

18. Sybirna, A., Wong, F. C. K. & Surani, M. A. Genetic basis for primordial germ cells specification in mouse and human: Conserved and divergent roles of PRDM and SOX transcription factors. Curr. Top. Dev. Biol. 135, 35–89 (2019).

19. Kalkan, T. et al. Complementary Activity of ETV5, RBPJ, and TCF3 Drives Formative Transition from Naive Pluripotency. Cell Stem Cell 24, 785–801.e7 (2019).

20. Zhang, J. et al. OTX2 restricts entry to the mouse germline. Nature 562, 595–599 (2018).

21. Calo, E. & Wysocka, J. Modification of enhancer chromatin: what, how, and why? Mol. Cell 49, 825–837 (2013).

22. Creyghton, M. P. et al. Histone H3K27ac separates active from poised enhancers and predicts developmental state. Proc. Natl. Acad. Sci. U. S. A. 107, 21931–21936 (2010).

23. Lara-Astiaso, D. et al. Immunogenetics. Chromatin state dynamics during blood formation. Science 345, 943–949 (2014).

24. Lee, K. et al. FOXA2 Is Required for Enhancer Priming during Pancreatic Differentiation. Cell Rep. 28, 382–393.e7 (2019).

25. Wang, A. et al. Epigenetic priming of enhancers predicts developmental competence of hESC-derived endodermal lineage intermediates. Cell Stem Cell 16, 386–399 (2015).

26. Lai, B. et al. MLL3/MLL4 are required for CBP/p300 binding on enhancers and super-enhancer formation in brown adipogenesis. Nucleic Acids Res. 45, 6388–6403 (2017).

27. Ostuni, R. et al. Latent enhancers activated by stimulation in differentiated cells. Cell 152, 157–171 (2013).

28. Sze, C. C. & Shilatifard, A. MLL3/MLL4/COMPASS Family on Epigenetic Regulation of Enhancer Function and Cancer. Cold Spring Harb. Perspect. Med. 6, (2016).

29. Bochyńska, A., Lüscher-Firzlaff, J. & Lüscher, B. Modes of Interaction of KMT2 Histone H3 Lysine 4 Methyltransferase/COMPASS Complexes with Chromatin. Cells 7, (2018).

30. Lin-Shiao, E. et al. KMT2D regulates p63 target enhancers to coordinate epithelial homeostasis. Genes Dev. 32, 181–193 (2018).

31. Wang, C. et al. Enhancer priming by H3K4 methyltransferase MLL4 controls cell fate transition. Proc. Natl. Acad. Sci. U. S. A. 113, 11871–11876 (2016).

32. Yan, J. et al. Histone H3 lysine 4 monomethylation modulates long-range chromatin interactions at enhancers. Cell Res. 28, 387 (2018).

33. Jang, Y. et al. H3.3K4M destabilizes enhancer H3K4 methyltransferases MLL3/MLL4 and impairs adipose tissue development. Nucleic Acids Res. 47, 607–620 (2019).

34. Placek, K. et al. MLL4 prepares the enhancer landscape for Foxp3 induction via chromatin looping. Nat. Immunol. 18, 1035–1045 (2017).

35. Ang, S.-Y. et al. KMT2D regulates specific programs in heart development via histone H3 lysine 4 di-methylation. Development 143, 810–821 (2016).

36. Ortega-Molina, A. et al. The histone lysine methyltransferase KMT2D sustains a gene expression program that represses B cell lymphoma development. Nat. Med. 21, 1199–1208 (2015).

37. Zhang, J. et al. Disruption of KMT2D perturbs germinal center B cell development and promotes lymphomagenesis. Nat. Med. 21, 1190–1198 (2015).

38. Dorighi, K. M. et al. Mll3 and Mll4 Facilitate Enhancer RNA Synthesis and Transcription from Promoters Independently of H3K4 Monomethylation. Mol. Cell 66, 568–576.e4 (2017).

39. Rickels, R. et al. Histone H3K4 monomethylation catalyzed by Trr and mammalian COMPASS-like proteins at enhancers is dispensable for development and viability. Nat. Genet. 49, 1647–1653 (2017).

40. Local, A. et al. Identification of H3K4me1-associated proteins at mammalian enhancers. Nat. Genet. 50, 73–82 (2018).

41. Rada-Iglesias, A. Is H3K4me1 at enhancers correlative or causative? Nature genetics vol. 50 4–5 (2018).

42. Ibarra-Soria, X. et al. Defining murine organogenesis at single-cell resolution reveals a role for the leukotriene pathway in regulating blood progenitor formation. Nat. Cell Biol. 20, 127–134 (2018).

43. Argelaguet, R. et al. Multi-omics profiling of mouse gastrulation at single-cell resolution. Nature (2019) doi:10.1038/s41586-019-1825-8.

44. Mohammed, H. et al. Single-Cell Landscape of Transcriptional Heterogeneity and Cell Fate Decisions during Mouse Early Gastrulation. Cell Rep. 20, 1215–1228 (2017).

45. Zylicz, J. J. et al. Chromatin dynamics and the role of G9a in gene regulation and enhancer silencing during early mouse development. Elife 4, (2015).

46. Tischler, J. et al. Metabolic regulation of pluripotency and germ cell fate through α-ketoglutarate. The EMBO Journal vol. 38 (2019).

47. Atlasi, Y. et al. Epigenetic modulation of a hardwired 3D chromatin landscape in two naive states of pluripotency. Nat. Cell Biol. 21, 568–578 (2019).

48. Hnisz, D. et al. Convergence of developmental and oncogenic signaling pathways at transcriptional super-enhancers. Mol. Cell 58, 362–370 (2015).

49. Yamaji, M. et al. Critical function of Prdm14 for the establishment of the germ cell lineage in mice. Nat. Genet. 40, 1016–1022 (2008).

50. Heintzman, N. D. et al. Distinct and predictive chromatin signatures of transcriptional promoters and enhancers in the human genome. Nat. Genet. 39, 311–318 (2007).

51. Rose, N. R. & Klose, R. J. Understanding the relationship between DNA methylation and histone lysine methylation. Biochim. Biophys. Acta 1839, 1362–1372 (2014).

52. Ooi, S. K. T. et al. DNMT3L connects unmethylated lysine 4 of histone H3 to de novo methylation of DNA. Nature vol. 448 714–717 (2007).

53. Buecker, C. et al. Reorganization of enhancer patterns in transition from naive to primed pluripotency. Cell Stem Cell 14, 838–853 (2014).

54. Magnúsdóttir, E. et al. A tripartite transcription factor network regulates primordial germ cell specification in mice. Nat. Cell Biol. 15, 905–915 (2013).

55. Hughes, A. L., Kelley, J. R. & Klose, R. J. Understanding the interplay between CpG island-associated gene promoters and H3K4 methylation. Biochim. Biophys. Acta Gene Regul. Mech. 1863, 194567 (2020).

56. Guo, X. et al. Structural insight into autoinhibition and histone H3-induced activation of DNMT3A. Nature 517, 640–644 (2015).

57. Zhang, Y. et al. Chromatin methylation activity of Dnmt3a and Dnmt3a/3L is guided by interaction of the ADD domain with the histone H3 tail. Nucleic Acids Res. 38, 4246–4253 (2010).

58. Li, B.-Z. et al. Histone tails regulate DNA methylation by allosterically activating de novo methyltransferase. Cell Res. 21, 1172–1181 (2011).

59. Nady, N. et al. Recognition of multivalent histone states associated with heterochromatin by UHRF1 protein. J. Biol. Chem. 286, 24300–24311 (2011).

60. Skvortsova, K. et al. Retention of paternal DNA methylome in the developing zebrafish germline. Nat. Commun. 10, 3054 (2019).

61. Hu, D. et al. The MLL3/MLL4 branches of the COMPASS family function as major histone H3K4 monomethylases at enhancers. Mol. Cell. Biol. 33, 4745–4754 (2013).

62. Xie, G. et al. MLL3/MLL4 methyltransferase activities regulate embryonic stem cell differentiation independent of enhancer H3K4me1. bioRxiv 2020.09.14.296558 (2020).

63. Li, J. et al. Accurate annotation of accessible chromatin in mouse and human primordial germ cells. Cell Res. 28, 1077–1089 (2018).

64. Zhang, Y. et al. Dynamic epigenomic landscapes during early lineage specification in mouse embryos. Nat. Genet. 50, 96–105 (2018).

65. Rulands, S. et al. Genome-Scale Oscillations in DNA Methylation during Exit from Pluripotency. Cell Syst 7, 63–76.e12 (2018).

66. Hui, T. et al. High-Resolution Single-Cell DNA Methylation Measurements Reveal Epigenetically Distinct Hematopoietic Stem Cell Subpopulations. Stem Cell Reports vol. 11 578–592 (2018).

67. Aubert, Y., Egolf, S. & Capell, B. C. The Unexpected Noncatalytic Roles of Histone Modifiers in Development and Disease. Trends Genet. 35, 645–657 (2019).

68. AlAbdi, L. et al. Oct4-Mediated Inhibition of Lsd1 Activity Promotes the Active and Primed State of Pluripotency Enhancers. Cell Rep. 30, 1478–1490.e6 (2020).

69. Maltby, V. E. et al. Histone H3K4 demethylation is negatively regulated by histone H3 acetylation in Saccharomyces cerevisiae. Proc. Natl. Acad. Sci. U. S. A. 109, 18505–18510 (2012).

70. Deshmukh, S., Ponnaluri, V. C., Dai, N., Pradhan, S. & Deobagkar, D. Levels of DNA cytosine methylation in the genome. PeerJ 6, e5119 (2018).

71. Barnett, K. R. et al. ATAC-Me Captures Prolonged DNA Methylation of Dynamic Chromatin Accessibility Loci during Cell Fate Transitions. Mol. Cell 77, 1350–1364.e6 (2020).

72. Sapozhnikov, D. M. & Szyf, M. Unraveling the functional role of DNA methylation using targeted DNA demethylation by steric blockage of DNA methyltransferase with CRISPR/dCas9. bioRxiv 2020.03.28.012518 (2020).

73. Ford, E. et al. Frequent lack of repressive capacity of promoter DNA methylation identified through genome-wide epigenomic manipulation. bioRxiv 170506 (2017).

74. Cruz-Molina, S. et al. PRC2 Facilitates the Regulatory Topology Required for Poised Enhancer Function during Pluripotent Stem Cell Differentiation. Cell Stem Cell 20, 689–705.e9 (2017).

75. Rauch, A. et al. Author Correction: Osteogenesis depends on commissioning of a network of stem cell transcription factors that act as repressors of adipogenesis. Nat. Genet. 51, 766 (2019).

76. Kaufman, C. K. et al. A zebrafish melanoma model reveals emergence of neural crest identity during melanoma initiation. Science 351, aad2197 (2016).

77. Pomerantz, M. M. et al. Prostate cancer reactivates developmental epigenomic programs during metastatic progression. Nat. Genet. 52, 790–799 (2020).

78. Acampora, D., Di Giovannantonio, L. G. & Simeone, A. Otx2 is an intrinsic determinant of the embryonic stem cell state and is required for transition to a stable epiblast stem cell condition. Development 140, 43–55 (2013).

79. Hayashi, K. & Saitou, M. Generation of eggs from mouse embryonic stem cells and induced pluripotent stem cells. Nat. Protoc. 8, 1513–1524 (2013).

80. Guo, G. et al. Klf4 reverts developmentally programmed restriction of ground state pluripotency. Development 136, 1063–1069 (2009).

81. Cong, L. et al. Multiplex genome engineering using CRISPR/Cas systems. Science 339, 819–823 (2013).

82. Calo, E. et al. Tissue-selective effects of nucleolar stress and rDNA damage in developmental disorders. Nature 554, 112–117 (2018).

83. Buenrostro, J. D., Giresi, P. G., Zaba, L. C., Chang, H. Y. & Greenleaf, W. J. Transposition of native chromatin for fast and sensitive epigenomic profiling of open chromatin, DNA-binding proteins and nucleosome position. Nat. Methods 10, 1213–1218 (2013).

84. Schmidl, C., Rendeiro, A. F., Sheffield, N. C. & Bock, C. ChIPmentation: fast, robust, low-input ChIP-seq for histones and transcription factors. Nat. Methods 12, 963–965 (2015).

85. Thomson, J. P. et al. CpG islands influence chromatin structure via the CpG-binding protein Cfp1. Nature 464, 1082–1086 (2010).

86. Clark, S. J. et al. Genome-wide base-resolution mapping of DNA methylation in single cells using single-cell bisulfite sequencing (scBS-seq). Nat. Protoc. 12, 534–547 (2017).

87. Zheng, G. X. Y. et al. Massively parallel digital transcriptional profiling of single cells. Nat. Commun. 8, 14049 (2017).

88. Trapnell, C. et al. The dynamics and regulators of cell fate decisions are revealed by pseudotemporal ordering of single cells. Nat. Biotechnol. 32, 381–386 (2014).

89. Nakaki, F. et al. Induction of mouse germ-cell fate by transcription factors in vitro. Nature 501, 222–226 (2013).

90. Stuart, T. et al. Comprehensive Integration of Single-Cell Data. Cell 177, 1888–1902.e21 (2019).

91. Bray, N. L., Pimentel, H., Melsted, P. & Pachter, L. Near-optimal probabilistic RNA-seq quantification. Nat. Biotechnol. 34, 525–527 (2016).

92. Melsted, P. et al. Modular and efficient pre-processing of single-cell RNA-seq. bioRxiv 673285 (2019).

93. La Manno, G. et al. RNA velocity of single cells. Nature 560, 494–498 (2018).

94. Li, H. & Durbin, R. Fast and accurate short read alignment with Burrows-Wheeler transform. Bioinformatics 25, 1754–1760 (2009).

95. Ramírez, F. et al. deepTools2: a next generation web server for deep-sequencing data analysis. Nucleic Acids Res. 44, W160–5 (2016).

96. Zhang, Y. et al. Model-based analysis of ChIP-Seq (MACS). Genome Biol. 9, R137 (2008).

97. McLean, C. Y. et al. GREAT improves functional interpretation of cis-regulatory regions. Nature Biotechnology vol. 28 495–501 (2010).

98. Rohde, C., Zhang, Y., Reinhardt, R. & Jeltsch, A. BISMA - Fast and accurate bisulfite sequencing data analysis of individual clones from unique and repetitive sequences. BMC Bioinformatics vol. 11 230 (2010).

99. Krueger, F. & Andrews, S. R. Bismark: a flexible aligner and methylation caller for Bisulfite-Seq applications. Bioinformatics vol. 27 1571–1572 (2011).

100. Martin, M. Cutadapt removes adapter sequences from high-throughput sequencing reads. EMBnet.journal vol. 17 10 (2011).

101. Langmead, B. & Salzberg, S. L. Fast gapped-read alignment with Bowtie 2. Nat. Methods 9, 357–359 (2012).

102. Clark, S. J. et al. scNMT-seq enables joint profiling of chromatin accessibility DNA methylation and transcription in single cells. Nat. Commun. 9, 781 (2018).

103. Yang, P. et al. Multi-omic Profiling Reveals Dynamics of the Phased Progression of Pluripotency. Cell Syst 8, 427–445.e10 (2019).

104. Dobin, A. & Gingeras, T. R. Mapping RNA-seq Reads with STAR. Curr. Protoc. Bioinformatics 51, 11.14.1–11.14.19 (2015).

105. Liao, Y., Smyth, G. K. & Shi, W. featureCounts: an efficient general purpose program for assigning sequence reads to genomic features. Bioinformatics 30, 923–930 (2014).

106. Anders, S. & Huber, W. Differential expression analysis for sequence count data. Nature Precedings (2010) doi:10.1038/npre.2010.4282.1.

107. Yi, L., Pimentel, H., Bray, N. L. & Pachter, L. Gene-level differential analysis at transcript-level resolution. Genome Biol. 19, 53 (2018).

